# Genetic mapping of *Citrullus amarus* for resistance to gummy stem blight in watermelon

**DOI:** 10.64898/2026.09.01.748752

**Authors:** Sourav Mahapatra, KB Kowsalya, Saheb Pal, Siddharood Maragal, Kuldeep Kumar, DC Lakshmana Reddy, Eguru Sreenivasa Rao

**Affiliations:** Division of Vegetable Crops, ICAR-Indian Institute of Horticultural Research, Bengaluru, India-560089; School of Crop Sciences, ICAR-Indian Agricultural Research Institute, Hazaribagh, Jharkhand, India-825405; Department of Vegetable Science, College of Horticulture and Forestry, CAU(I), Pasighat, Arunachal Pradesh, India-791102; ICAR- Indian Institute of Pulses Research, Kanpur, Uttar Pradesh, India-208024; Division of Basic Sciences, ICAR-Indian Institute of Horticultural Research, Bengaluru, India-560089

**Keywords:** Biotic stress, Disease, Genetics, Inheritance, Mapping, QTL-seq

## Abstract

Gummy stem blight affects several cucurbit crops and has emerged as a serious production constraint of watermelon cultivation globally. However, no commercial varieties are available possessing resistance against this disease. Identification of genetic determinants and underlying QTLs are prerequisites for resistant variety development. In this direction, an attempt was made to understand genetics and identify responsible loci using two resistant accessions *viz*., PI 482276 and IIHR 82. Inheritance analysis indicated that resistance to GSB is governed by polygenes in both accessions. Four QTLs were identified for the population derived from PI 482276 × Arka Manik, while six were observed in the cross IIHR 82 × Arka Manik employing QTL-seq approach. The QTL regions contained important genes putatively involved in resistance reaction such as NAC domain-containing protein 17-like, chitinase 2- like, Pectinesterase, Coiled-coil domain-containing protein 18, Kinase family protein expansin-like B1 protein etc. Our results offer further insight into the disease resistance genetics and its utilization in resistance breeding for gummy stem blight in watermelon.

## Introduction

Watermelon, *C. lanatus* (Thunb.) Matsum. & Nakai, is the most cultivated cucurbit crop both in area and production. The global area under watermelon cultivation is around 3.05 million hectares, producing 101.62 million tonnes (FAO 2022). Despite being a commercial crop in India, there is a huge gap between national and global productivity. Among the various biotic production constraints, Gummy Stem Blight (GSB), caused by the fungus *Didymella bryoniae* (Auersw.) Rehm, is perhaps the most destructive fungal disease, resulting in substantial economic loss (Keinath et al. 1995; Bruton 1998; Stewart et al. 2015). It has been reported in six continents to infect at least 12 genera and 23 species of Cucurbitaceae (Keinath 2011; Luo et al. 2022).

The Gummy Stem Blight disease is caused by three morphologically similar but genetically distinct species of *Stagonosporopsis viz*., *S. cucurbitacearum* (syn. *D. bryoniae*), *S. citrulli*, and *S. caricae* (Stewart et al. 2015), producing similar symptoms in plants. GSB occurs mostly during warm and humid environmental conditions that are congenial for disease development (Keinath et al. 1995; Robinson and Decker-Walters 1997; Keinath 2011; Babu et al. 2015; Stewart et al. 2015). The pathogen is a facultative necrotroph (Svedelius 1990) and enters into the plants, particularly through natural wounds of the older leaves (Blakeman 1971; Pharis et al. 1982; Van Steekelenburg 1985; Svedelius 1990). The pathogen damages different plant parts, including leaves, stems, flowers, fruits, tendrils, pedicels and seeds (Keinath 2013). Disease symptoms start with marginal brown discolouration of the leaves, spreading toward the inner leaf lamina. Later, the pathogen spreads to cover completely, turning it dark brown and dry, affecting photosynthesis, ultimately resulting in mortality and may cause complete yield loss (Mahapatra et al. 2020). The wide host range, seed, soil and air-borne nature of dispersal make the fungus difficult to eradicate with fungicides. Moreover, the rapid development of resistance towards fungicides leads to an environmental concern, making fungicidal management unsustainable (Ren et al. 2020). Therefore, the utilization of host resistance is considered to be a preferred alternative.

The disease has been reported in different countries across many cucurbits. However, to date, no source of resistance to GSB has been reported in the cultivated background of watermelon though several wild accessions of watermelon *viz*., PI 189225 (Sowell and Pointer 1962; Gusmini et al. 2005, 2017), PI 482276 (Gusmini et al. 2005), PI 271778 (Sowell 1975), PI 500335, PI 505590, PI 512373, PI 164247, and PI 500334 (Boyhan et al. 1994) have been reported as resistant. We have recently reported two watermelon accessions *viz*., IIHR 82 and PI 482276 to be resistant to an Indian isolate (NCBI accession no. KC460840.1) of *Stagonosporopsis cucurbitacearum* (Mahapatra et al. 2022). To effectively utilize these accessions, it is necessary to understand the genetics of resistance to plan a deployment strategy. Availability of the reference genome of watermelon and next-generation sequencing techniques can help to locate the regions containing genes or QTLs associated with resistance employing QTL-seq approach (Liu et al. 2012; Takagi et al. 2013). Thus, the present research work was undertaken to study the inheritance pattern and identify QTL regions in the two GSB-resistant accessions identified in Indian conditions employing QTL-seq approach.

## Materials and Methods

The present experiment was undertaken at ICAR-Indian Institute of Horticultural Research, Bengaluru, India during 2019-23. The experimental plot is located at 13°07′41.1″N latitude, 77°29′34.1″E longitude and 890 m above the mean sea level. The experimental material comprised two populations (F_2:3_) derived from the resistant accessions belonging to *Citrullus amarus viz*., IIHR 82 and PI 482276, which were crossed with susceptible parent, Arka Manik (*C. lanatus*). The resistant accessions showed resistance both under polyhouse and natural epiphytotic conditions (Mahapatra et al. 2022). The line, IIHR 82 is a red-seeded *C. amarus* accession being maintained at ICAR-IIHR; PI 482276 is a germplasm accession collected from the USDA; Arka Manik is a popular mega variety of watermelon developed at ICAR-IIHR and is susceptible to GSB. NS 295 is a popular commercial variety widely cultivated in this region, developed by Namdhari Seeds Pvt. Ltd., Karnataka, India and was used as a commercial check during the field evaluation. Polyhouse and field screening of mapping populations was carried out to evaluate the genetics of resistance to GSB.

## Development of populations for genetic analysis

During spring 2019, resistant parents (PI 482276 and IIHR 82) were crossed as female parents, with Arka Manik, the susceptible parent as male. The mature seeds were harvested from the crossed fruits and the F_1_ seeds were sown in the rainy season 2020 were self-pollinated to generate F_2_ and backcrossed to both resistant (P1) and susceptible parents (P2) to develop BCP1F_1_ and BCP2F_1_ populations, respectively. During the *rabi* season of 2020 self-pollination was done in each F_2_ plant to obtain F_3_ seeds and in the backcrossed plants to derive the BCP1F_1:2_ and BCP2F_1:2_ seeds for phenotyping. The fruit morphological features of the parental accessions, the F_1_ and the F_2_ progenies, have been depicted in Figure 1.

**Figure 1.**
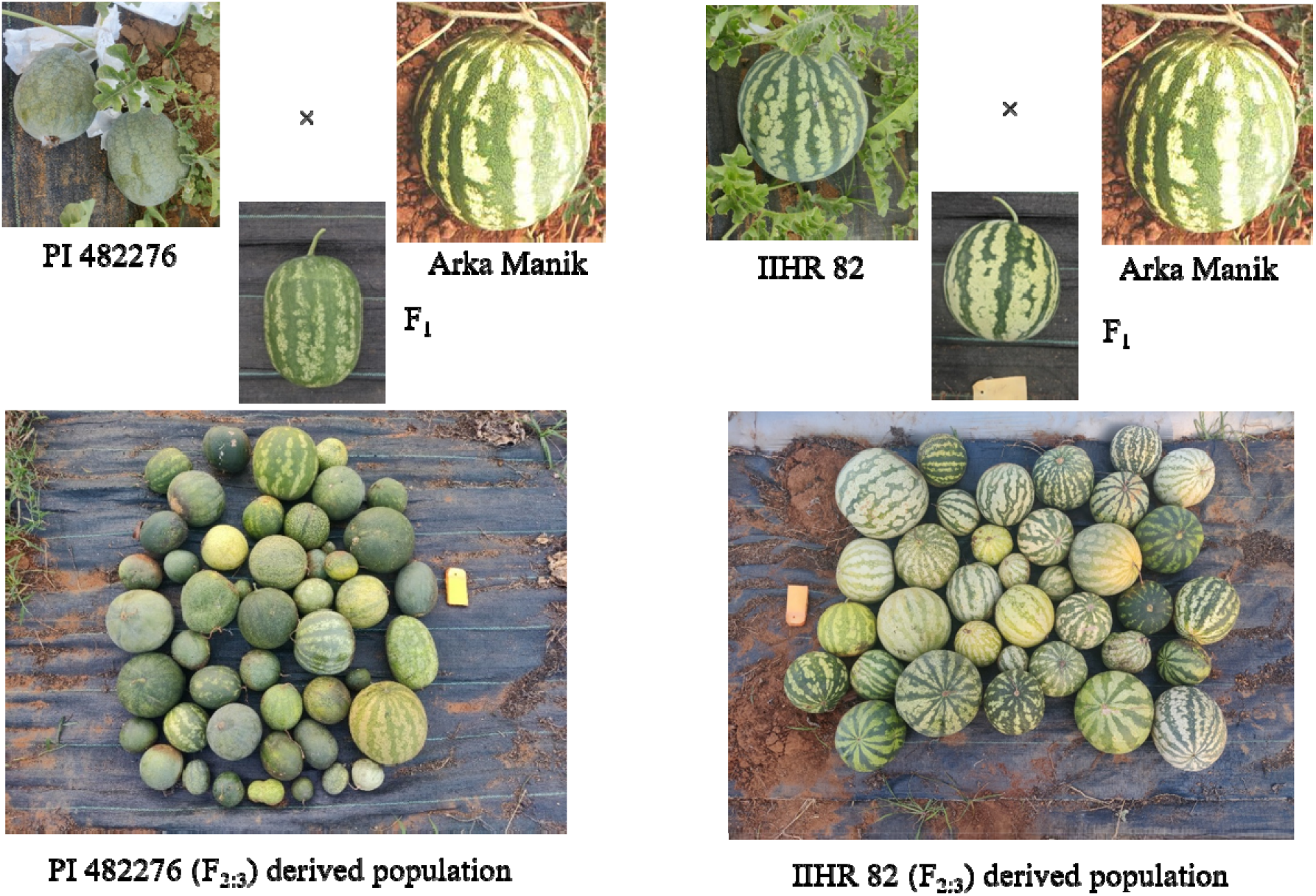
The morphological features of the parental accessions, their F_1_ and F_2:3_ progenies

## Screening of populations employing artificial inoculation

During the *rabi* season of 2021, these populations (F_1_, BCP1F_1:2_, BCP2F_1:2_, and F_3_) were sown along with resistant and susceptible parents and susceptible checks in a polyhouse to study their reaction toward the pathogen. The genotypes were sown in 98 celled portrays and 14 seedlings/genotype were raised in two replications of seven plants each. The mycelial disc inoculation was followed to challenge the plants with the virulent isolate KC460840.1. The screening protocol, as described by Mahapatra et al. (2020), was followed. Observations were recorded starting from seven days of inoculation and repeated at seven-day intervals till 21 days after inoculation. The scale of Gusmini et al. (2002) was adopted with suitable modification for quantifying the disease reaction. The rating 0- for only restricted leaf infection; 1- for leaf and petiole infection; 2- for leaf, petiole and stem infection; 3- for complete drooping of the plant. Disease ratings and analysis of PDI and AUDPC are performed as described in Mahapatra et al. (2020).

## Screening of populations in a sick plot

During the *rabi* season of 2021, the P_1_, P_2_, F_1_ and F_2:3_ populations of both the crosses were sown in portrays and 14-day seedlings were transplanted in a GSB-infected sick plot following randomized block design with two replications of seven plants each. Paired row planting on raised beds was followed with entries on one side and susceptible check (NS 295) on the other side. No fungicides were sprayed throughout the screening to allow symptom development.

Observations were recorded starting from 50 days after planting (DAP). Further observations were recorded at 10-day intervals until 90 DAP when the maximum number of plants of the susceptible parent and susceptible check completely died. A scale given by Dos-Santos et al. (2016) was suitably modified and used for this evaluation; where 0– for no symptoms, 1– for drooping, wilting and blight of the leaves at the collar region, 2– for drooping, wilting and blight of the leaves up to half of the vine, 3– for severe wilt of plant and 4– for complete mortality of the plant. The percent disease index (PDI) was calculated based on scoring data of individual seedlings using the formula of McKinney (1923):

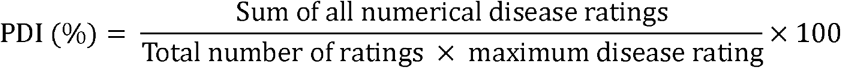

## Six-generation mean analysis

At 21 DPI, the PDI values observed in artificial screening were used to study the genetics and gene action for GSB resistance. PDI values were subjected to a normality test before being grouped into resistant or susceptible classes. The disease reaction of individual plants in terms of PDI was subjected to a scaling test (Mather 1949; Hayman and Mather 1955), joint scaling test (Cavalli 1952) and generation mean analysis employing six-parameter model (Hayman 1958; Jinks and Jones 1958). Data analysis was carried out using SAS v. 9.3 software.

The following Castle□Wright’s formula (Castle 1921) was employed to compute the minimum number of effective gene blocks involved in resistance: (P1−P2)^2^ /8(VF_2_−VF_1_) where, n=minimum number of effective gene blocks involved in resistance, P1 and P2= parental means, VF_1_ and VF_2_ are variances in the disease reaction of first and second filial generations, respectively.

## QTL-seq analysis

The population for QTL analysis consisted of 55 F_2_ individuals derived from PI 482276 × Arka Manik and 76 F_2_ from IIHR 82 × Arka Manik. Individual plants were tagged and DNA was isolated from young leaves using the CTAB method (Doyle and Doyle 1987). The most resistant and susceptible families were identified based on the disease screening data from both polyhouse and field evaluations. F_2_ individual plant classifications were made into resistant and susceptible categories based on the disease reaction of corresponding F_2:3_ families. Six F_2:3_ extreme families from each category were identified, and their corresponding F_2_ plant DNA was pooled in equal quantities to construct the resistant and susceptible bulks for both populations. DNA was quantified using a Denovix DX-11 spectrophotometer (DeNovix Inc., USA) and normalized to a concentration of 1058.62 ng/µl. The parental DNA with resistant and susceptible bulks were subjected to whole-genome sequencing-based Bulked Segregant Analysis (BSA) using Illumina Hi-Seq 2500 platform using NEB adapters to generate 150 base pair long paired-end reads at Sandor Life Sciences Private Limited, Hyderabad, India.

The high quality reads were subjected to FastQC (Andrews 2010) for quality check. The poor quality reads and adaptors were removed using trimmomatic v0.39 (Bolger et al., 2014) to obtain high quality fastq sequences for three samples in each cross, viz., resistant parents (PI 482276 and IIHR 82), resistant bulk and susceptible bulk. The quality reads of resistant parents (PI 482276 and IIHR 82) were aligned with the available reference genome of elite Chinese watermelon line 97103 v.2 (Guo et al. 2013) using BWA v. 0.5.9 (Li and Durbin 2009). By replacing the alleles of cv. 97103 with that of the SNPs from the resistant genotypes, a genomic FASTA file for the resistant parent was imputed following the procedure suggested by Takagi et al. (2013). High-quality reads of the bulks (both resistant and susceptible) were aligned to the imputed reference FASTA using BWA v. 0.5.9 (Li and Durbin 2009). The variants were filtered through Coval v.1.4.1 at a mutation index of 4 and read depth threshold of 7 to improve the accuracy of SNP calling (Kosugi et al. 2013). Conversion of alignments to SAM/BAM files was carried out using SAMtools v. 0.1.8 (Li et al. 2009). The ΔSNP index was calculated based on the read depth (Takagi et al. 2013) using the QTLseq pipeline. The results were plotted using a 1 MB window size and 10 kb increment.

## Results

### Screening of the genetic populations for reaction to GSB

The results of the disease development (21 DPI) in artificial inoculation under polyhouse for all six generations and in a sick plot under natural epiphytotic conditions for F_2:3_ families (75 DAP) along with resistant and susceptible parents have been presented in Supplementary Tables 1 and 2. The number of plants screened has been presented in Supplementary Table 3. Under artificial inoculation, the F_2:3_ population derived from PI 482276 × Arka Manik showed a PDI range of 50% to 100%. The mean disease score at 21 DPI for the resistant parent was 53.33%, whereas for the susceptible parent was 91.41% (Figure 2A). Further, the mean PDI for the F_1_, BC1F_1_ and BC2F_1_ populations were 61.64%, 65.96% and 75.28%, respectively at 21 DPI. The PDI for the F_2:3_ population derived from IIHR 82 × Arka Manik ranged from 48.89% to 100.00%. The mean PDI for the resistant parent was 63.31%, whereas for the susceptible parent was 94.18% (Figure 2B). Further, the mean disease reaction for the F_1_, BCP1F_1_ and BCP2F_1_ populations were 66.06%, 72.14% and 80.53%, respectively at 21 DPI.

**Figure 2.**
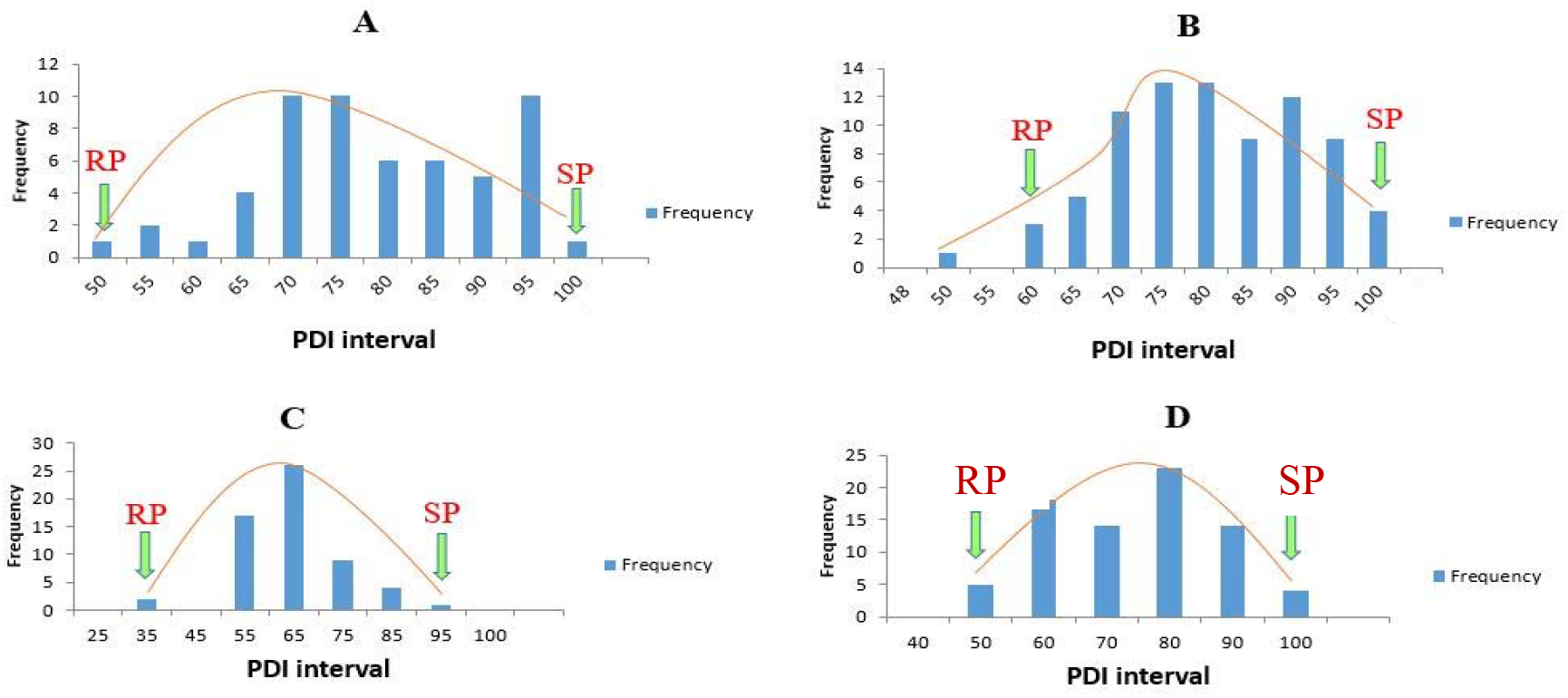
Graphical representation of PDI values of parents and families screened under artificial polyhouse (A and B) conditions and natural epiphytotic conditions in field (C and D) against GSB. A and C: F_2:3_ families derived from PI 482276 and B and D: F_2:3_ families derived from IIHR 82.

Natural epiphytotic screening for the F_2:3_ families derived from PI 482276 × Arka Manik showed a PDI range of 47.5 to 84.65. The resistant parent PI 482276 recorded a PDI of 27.82%, whereas the susceptible parent, Arka Manik showed 91.67% (Figure 2C). Field screening for the F_2:3_ families derived from IIHR 82× Arka Manik showed a PDI range of 47.50 to 92.50. The resistant parent, IIHR 82 recorded a PDI of 44.59%, whereas the susceptible parent Arka Manik showed 91.67% at 75 DAP (Figure 2D). The susceptible check NS 295, planted as a paired row along the entries, recorded 100% PDI, indicating uniform inoculum exposure among all entries tested.

### Six-generation mean analysis

The PDI values of the generations obtained during screening under the polyhouse were subjected to a normality test (P=0.005) for skewness and kurtosis. Results for the cross PI 482276×Arka Manik showed a negative skewness of -0.029 with a kurtosis of -0.538. The population IIHR 82×Arka Manik recorded a skewness of -0.188 with a kurtosis of -0.461 (Table 1). For both populations, skewness is near ‘0’, making the distribution of their PDI nearly normal or continuous. In both crosses, the distribution of F_2:3_ populations showed negative skewness and negative kurtosis (Table 1).

**Table 1.** Distribution analysis of PDI of the progenies derived from PI 482276 × Arka Manik and IIHR 82 × Arka Manik.

| Parameter | PI 482276 × Arka Manik | IIHR 82 × Arka Manik |
| --- | --- | --- |
| Mean | 76.7637 | 78.22436 |
| Standard Error | 1.568551 | 1.214365 |
| Median | 75.005 | 78.245 |
| Mode | 73.34 | 66.67 |
| Standard Deviation | 11.52645 | 10.72498 |
| Sample Variance | 132.8591 | 115.0253 |
| Kurtosis | -0.53878 | -0.46181 |
| Skewness | -0.02926 | -0.18819 |
| Range | 50 | 51.11 |
| Minimum | 50 | 48.89 |
| Maximum | 100 | 100 |
| Sum | 4145.24 | 6101.5 |

The PDI values of all six generations in both crosses were subjected to a scaling test (Table 2). In PI 482276×Arka Manik, the components A, C and D were significant, whereas, in IIHR 82×Arka Manik, only A and C components were significant. The significance of the D component in the former implied the predominance of additive × additive type of epistasis, while the significance of the C component of epistasis in the latter revealed the prevalence of dominance×dominance type epistasis. This confirms the polygenic nature of resistance through nonallelic interactions.

**Table 2.**
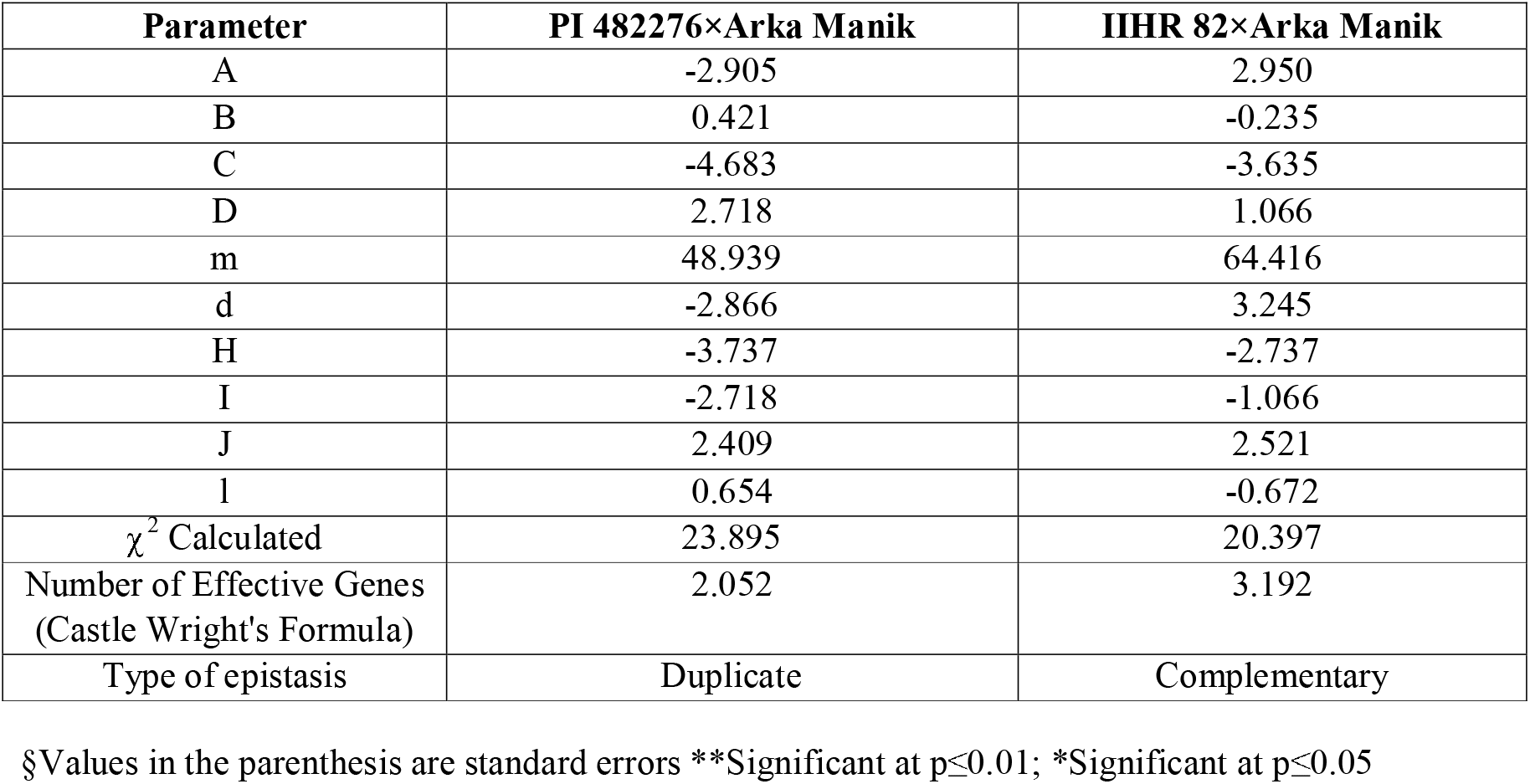
Scaling test and gene effects for GSB resistance as assessed in two resistant × susceptible crosses using a six-parameter model.

Different genetic components *viz*., mean (m), additive (d), dominance (h), additive × additive (i), additive×dominance (j) and dominance×dominance (l) effects for both the crosses were assessed and presented in Table 2. In the cross PI 482276×Arka Manik, mean (48.939), additive (-2.866) and dominance components (-3.737) were significant along with additive×additive (-2.718) and additive×dominance (2.409) type of epistasis. It showed the prevalence of both additive and non-additive interactions in this combination. The contrasting signs of dominance (h) and dominance×dominance (l) suggested the epistasis to be of duplicate type. The population derived from IIHR 82×Arka Manik recorded significant mean (64.416), additive (3.245) and dominance components (2.737) along with additive×dominance (2.521) type of epistasis. Similar signs of dominance (h) and dominance×dominance (l) suggested complementary type epistasis. The number of effective genes for the cross PI 482276×Arka Manik was 2.052 and for the cross IIHR 82×Arka Manik was 3.192, calculated using the Castle-Wright’s formula.

### Quantitative trait loci governing resistance to GSB

The whole genome sequencing of the resistant bulk, susceptible bulk and resistant parent for the population PI 482276×Arka Manik generated 9.41 GB, 7.86 GB and 6.75 GB data, respectively. IIHR 82×Arka Manik generated 6.69 GB, 7.06 GB and 7.85 GB of data, respectively. The poor-quality reads were filtered and the quality reads (>Q30) comprising 93.04 %, 93.25 % and 92.92 % of the total reads of resistant parent, resistant bulk and susceptible bulk of PI 482276×Arka Manik population were considered for further analysis. Similarly, for the population, IIHR 82×Arka Manik quality reads (>Q30) comprising 92.93%, 93.17% and 93.14% of the total reads of resistant parents, resistant bulk and susceptible bulk were used for further analysis. Across all chromosomes, 108157 SNPs and 43322 InDels were identified for PI 482276. The highest number of SNPs and InDels were identified in chromosome 5 (n=13744 and 5062, respectively). While in IIHR 82, the highest number of SNPs (1754) and InDels (1288) were identified in chromosome 9.

QTL-seq recorded four significant QTLs (at confidence interval 95) each on chromosomes 1, 3, 6 and 8 for GSB resistance in PI 482276×Arka Manik (Figure 3).

**Figure 3.**
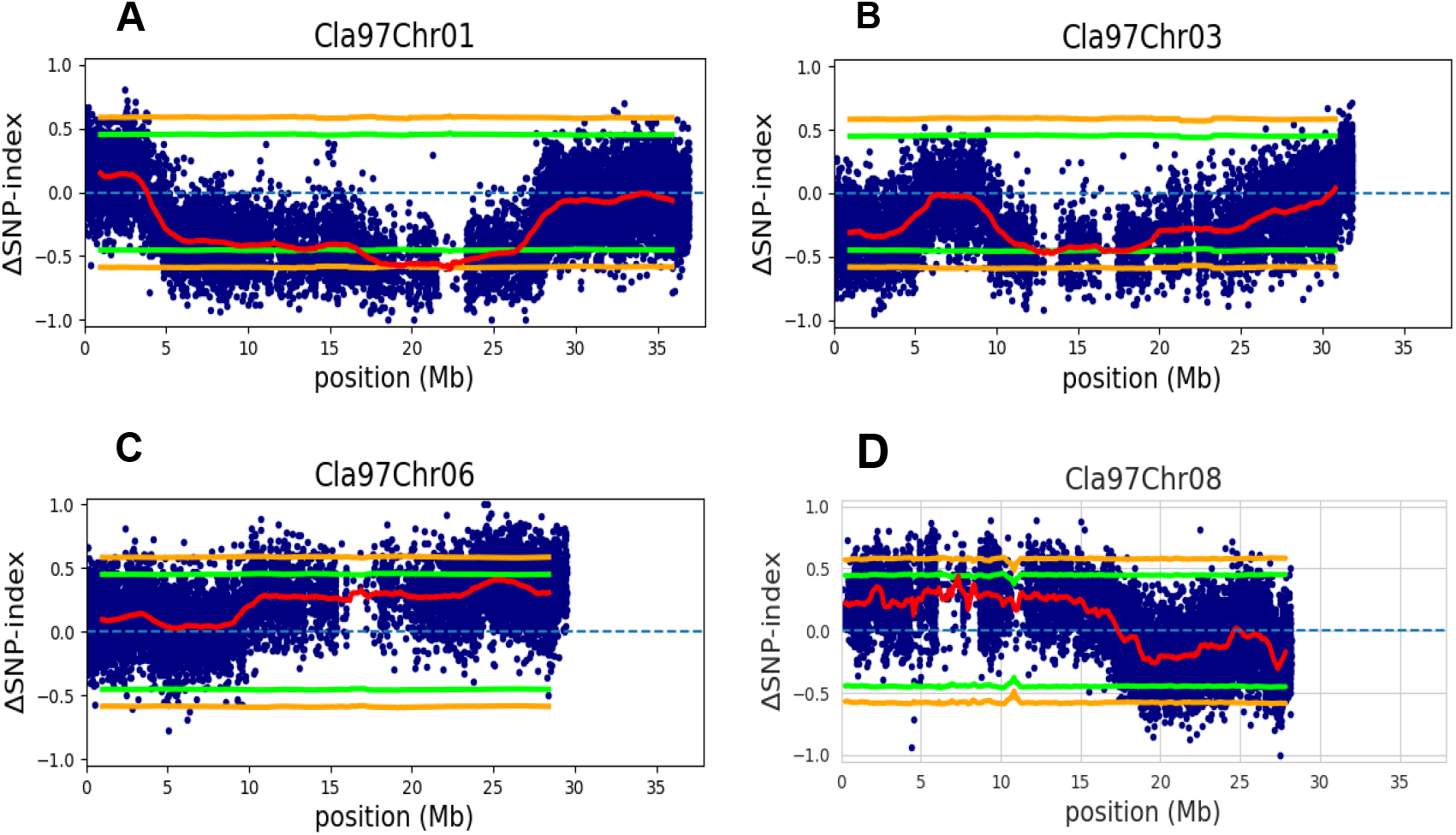
Statistically significant QTLs identified for GSB resistance in PI 482276 × Arka Manik (Green line: Confidence Interval 95 and Orange line: Confidence Interval 99).

On Chromosome 1, the identified significant QTL region spanned from 17.74 Mb to 23.07 Mb, peaking at 22.29 Mb (Figure 3-A). The QTL region holds a total of 84 genes. Among them *Cla97C01G011680* (Cytochrome P450 family protein) which is involved in wound healing reactions and oxidase reductase activity, *Cla97C01G011810* (Stress-response ab barrel domain-containing protein), *Cla97C01G011820* (wall-associated receptor kinase 4- like), *Cla97C01G011380* (NF-kappa-B inhibitor-like protein 2 isoform 1), *Cla97C01G011510* (Peroxidase), *Cla97C01G011710* (histidine-containing phosphotransfer protein 5-like) performing signal transduction activity are the important putative genes involved in disease resistance reactions. On chromosome 3, the QTL region spanned from 11.80 Mb to 18.96 Mb with a peak between 12.50 to 13.54 Mb region (Figure 3-B). This region includes a total of nine genes. Likewise, on chromosome 6 a single QTL spanned from 24.65 Mb to 26.22 Mb with a peak at 25.34 Mb (Figure 3-C) containing 14 genes. Among them, the genes with putative resistance roles include *Cla97C06G111810* (Kinase-like protein) and *Cla97C06G111770* (Carotenoid cleavage dioxygenase) governing oxidase reductase activities. The QTL spans from 7.1 to 7.5 Mb for chromosome 8, with a peak observed at 7.3 Mb (Figure 3-D). This region contains a total of 10 genes. Among them, *Cla97C08G146850* (protein kinase family protein), *Cla97C08G146890* (E3 ubiquitin-protein ligase listerin-like), *Cla97C08G146900* and *Cla97C08G146930* (Cysteine protease, putative) are the probable major genes responsible for resistance reactions.

Six significant genomic regions were found for the population IIHR 82 × Arka Manik, one each on chromosomes 6 and 7 and two each on chromosomes 2 and 9, conferring resistance to GSB and presented in Figure 4.

**Figure 4.**
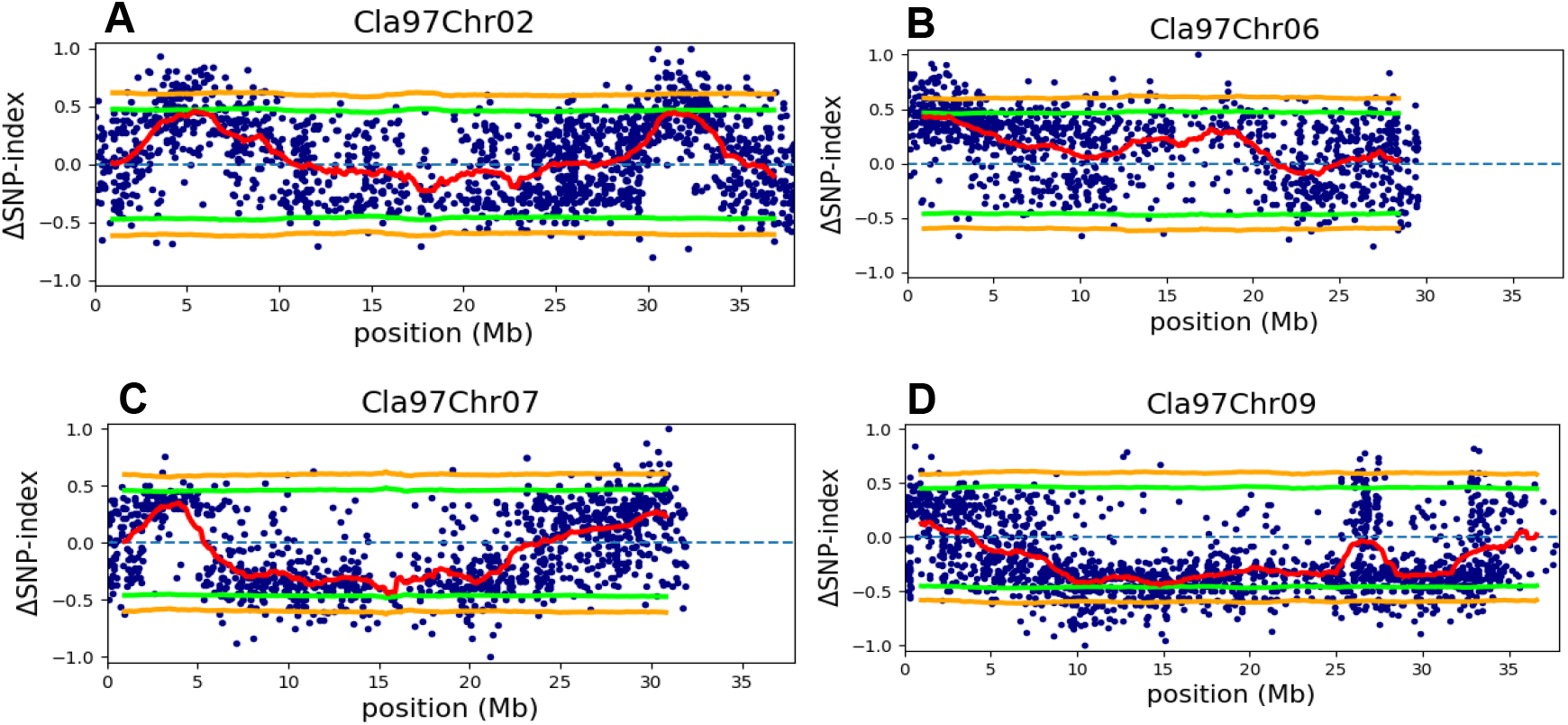
Statistically significant QTLs identified for GSB resistance in IIHR 82 × Arka Manik (Green line: Confidence Interval 95 and Orange line: Confidence Interval 99).

The first QTL on chromosome 2 spanned between 4.28 Mb to 6.42 Mb with a peak at 5.30 Mb (Figure 4-A). This genomic region contains a total of 29 genes. The putative genes involved in resistance reaction in this region are *Cla97C02G026800* (chitinase 2- like), *Cla97C02G026890* (protein MKS1-like) involved in resistance gene-dependent defence response signaling pathway, Cla97C02G026720 (Pectin lyase-like superfamily protein), *Cla97C02G026840* (Pectinesterase), *Cla97C02G026780* (Coiled-coil domain-containing protein 18, putative isoform 2). Similarly, the second QTL spanned between 30.67 Mb and 32.36 Mb, peaking at 31.35 Mb (Figure 4-A). The total number of genes present in this region was 23. The major putative genes governing resistance reaction are *Cla97C02G030120* (protein LAZ1 isoform X1) involved in hypersensitive response and programmed cell death, *Cla97C02G030150* (lysM domain-containing GPI-anchored protein 2) having chitin-binding activity, *Cla97C02G029960* (Alpha-1,3-mannosyl-glycoprotein beta-1,2-N-acetylglucosaminyltransferase), *Cla97C02G030040* (Kinase family protein) involved in cell surface receptor signaling pathway and *Cla97C02G030050* (Hexosyltransferase) performing polygalacturonate 4-alpha-galacturonosyltransferase activity. On chromosome 6, a single QTL spanned from 0.78 Mb to 2.7 Mb, peaking at 2.48 Mb (Figure 4-B). This region holds 177 genes, among which *Cla97C06G110780* (Cellulose synthase) *Cla97C06G111500* (calmodulin-binding transcription activator 4), *Cla97C06G111150* (hydroxyproline-rich glycoprotein family protein, putative), *Cla97C06G111140* (Leucine-rich repeat receptor-like protein kinase family), *Cla97C06G111130* (Leucine-rich repeat receptor-like kinase protein), *Cla97C06G110840* (Receptor kinase), *Cla97C06G110270* (Peroxidase) and *Cla97C06G110350* (Trigger factor) are the probable genes with resistance like function. On chromosome 7, the QTL started from 15.22 Mb to 16.01 Mb, peaking at 15.45 Mb (Figure 4-C). It contains three genes in the identified QTL region. The probable major gene involved in the resistance reaction is *Cla97C07G134690* (MLP-like protein 328), a defence response gene along with *Cla97C07G134710* (Cytochrome P450). Among the two QTLs on chromosome 9, the first spanned from 9.58 Mb to 11.39 Mb with a peak at 11.27 Mb, whereas the second QTL spanned from 13.87 Mb to 16.12 Mb with a peak at 14.66 Mb (Figure 4-D). The first QTL harbours a total of 152 genes. Among them, *Cla97C09G173270* (NAC domain-containing protein 7) *Cla97C09G173490* (NAC domain-containing protein 17-like), *Cla97C09G174440* (E3 ubiquitin-protein ligase RNF14) are the probable genes playing role in the resistance response. The second QTL holds a total of 39 genes. Among them, *Cla97C09G175430* (Casein kinase II subunit beta), *Cla97C09G175480* (expansin-like B1) and *Cla97C09G175580* (Ras-related GTP-binding protein) are important resistance-like genes.

## Discussion

### Studies on the inheritance of resistance to GSB in watermelon genotypes

Gummy stem blight is a major disease of watermelon globally (Gusmini et al. 2017). Management of this disease through cultural management and fungicidal spray is not completely effective and no resistance has been reported from a cultivated background. Thus, inheritance studies of GSB resistance are mostly confined to wild relatives of watermelon, particularly to *C. amarus*. A highly resistant wild watermelon genotype PI 189225 belongs to this species and is a frequently used accession for studying genetics as well as mapping the resistant loci (Norton et al. 1979; Gusmini et al. 2017; Ren et al. 2020).

Though several accessions are reportedly resistant to this disease, the inheritance of resistance is not well understood. Earlier reports demonstrated the presence of a single recessive gene ‘*db*’ in the cross between ‘PI 189225’ and ‘Charleston Grey’ (Norton et al. 1979). Other resistant accessions, such as PI 482283 and PI 526233 were also evaluated for understanding genetics (Gusmini et al. 2017). Efforts were made to develop variety by introducing the ‘*db*’ gene. However, when evaluated under natural epiphytotic conditions, the resultant cultivars possessed lesser resistance than the resistant parent. In crosses involving PI 189225 × NH Midget, PI 482283 × NH Midget, PI 482283 × Calhoun Gray and PI 526233 × All Sweet, the genetics of disease reaction was elusive. For all four cross combinations under field and polyhouse conditions except for PI 482283 × Calhoun Gray, the BC1F_1_ segregation ratio rejected the null hypothesis of a single gene controlling the GSB resistance. For PI 482283 × Calhoun Gray, the greenhouse tests did not support the field validation of a monogenic recessive gene controlling the resistance reaction to GSB (Gusmini et al. 2017).

In our experiments, we identified two genotypes *viz*., PI 482276 (previously reported by Gusmini et al. 2005) and IIHR 82 (a newly identified source) as resistant to Indian isolate of pathogen causing GSB in watermelon (Mahapatra et al. 2020, 2022). A clear phenotypic difference was observed between the resistant and susceptible parents, making them suitable for this inheritance study. We considered a common susceptible parent for a clear comparison between the genetics of resistance in both crosses, as any difference in performance shall be only contributed by the corresponding resistant parent. The PDI recorded from the polyhouse screening experiment of F_2:3_ families was subjected to a normality check. The skewness value was near zero for both populations and the kurtosis is negative. This suggests that the distribution is almost normal, showing a continuous phenotypic distribution of disease severity for both populations due to polygenic inheritance. The sign of kurtosis is negative (platykurtic) for both populations, indicating the involvement of many genes in the expression of resistance along with background modifiers. It also shows the presence of wider variability in the segregating population, offering an opportunity for selection.

The scaling test and the six-generation mean analysis confirmed the presence of non-allelic interaction or epistatic gene action for both crosses. Generation mean analysis showed the epistasis to be a complementary type in IIHR 82 × Arka Manik, suggesting that pedigree selection would be helpful to improve resistance in crosses considering IIHR 82 as a donor. As per Castle Wright’s Formula, the minimum number of effective genes is 3.192. In the cross, PI 482276 × Arka Manik, the involvement of both additive and non-additive genes with a duplicate type of epistasis interaction was observed. Hence, reciprocal recurrent selection followed by simple selection seems to be a more appropriate breeding scheme involving PI 482276 as a donor parent. Further, the minimum number of effective genes estimated is 2.052 in PI 482276.

In summary, the genetics of resistance to GSB was found to be polygenic, involving interaction effects as the classification of the segregating population into a discrete class (resistant/susceptible) was not possible due to a continuous variation. Further partial compatibility leading to unviable seeds in segregating progenies complicates the deployment of these resistance sources through traditional breeding methods. Therefore, to ease the transfer of resistance into a desirable background, we tried to identify the major QTLs governing resistance in both resistant accessions.

### Identification of QTLs conferring resistance

Among cucurbits, QTLs governing resistance to GSB have been reported initially in cucumber (Lou et al. 2013; Liu et al. 2017; Zhang et al. 2017). Lou et al. (2013) reported QTLs for GSB resistance on chromosomes 4 (*GSB4*) and 6 (*GSB6b*) with a span width of 12 cM and 11 cM respectively. Similarly, Liu et al. (2017) reported a total of six QTLs (on chromosomes 3, 4, 5 and 6), among which, the QTL on chromosome 5 (*gsb5*.*1*) was found to be stable with a 0.5 cM span width, harboring seven resistance genes. Zhang et al. (2017) identified five QTLs conferring resistance to GSB on cucumber. Among these, the QTL on chromosome 6 (*gsb-s6*.*2*) accounted for the highest phenotypic variation. Fourteen disease-resistance genes had been identified in that region spanning 3.2 cM.

In watermelon, PI 189225 and PI 482276 are two most studied accessions for mapping of GSB resistance. QTLs have recently been identified in PI 189225, *viz*., *Qgsb8*.*1*, a major QTL on the short arm of chromosome 8, explaining 32% phenotypic variation (Ren et al. 2020). For the same accession, two major QTLs, *qLL8*.*1* and *qSB8*.*1* on chromosome 8 and one minor QTL, *qSB6*.*1* on chromosome 6 were identified explaining 10.5%, 10.0% and 9.7% of phenotypic variation, respectively (Lee et al. 2021). Similarly, Adams et al. (2022) identified QTLs on chromosomes 2, 5, 9 and 11 in the cross Sugar Baby × PI 189225 (*Citrullus amarus*). They identified a novel QTL on chromosome 5 (*Qgsb5*.*2*) explaining 13% of phenotypic variation towards the resistance to *S. citrulli*. The other most studied resistant accession PI 482276, also used in the current study, was previously reported to carry 3 QTLs *i*.*e*., *ClGSB3*.*1, ClGSB5*.*1* and *ClGSB7*.*1* on chromosomes 3, 5 and 7, respectively explaining between 6.4% and 21.1% of phenotypic variation (Gimode et al. 2020). Recently, Hong et al. (2022) identified three QTLs namely, *ClGSB1*.*1, ClGSB10*.*1* and *ClGSB11*.*1* associated with GSB resistance in the cross PI 279461×PI 223764. Among them, the QTL on chromosome 1 (*ClGSB1*.*1*) is a major QTL holding five candidate genes and explaining about 10% of phenotypic variation.

In the current study, a total of 10 QTLs on 7 chromosomes (1, 2, 3, 6, 7, 8 and 9) have been identified. QTLs have earlier been reported on all these chromosomes (Table 3). Among them, eight QTLs are physically overlapping/adjoining those reported by earlier workers on chromosomes 1, 3 and 6 for PI 482276 and on chromosomes 2, 6 and 9 for IIHR-82. These might be considered as hotspots for further study. For the other two QTLs, though chromosomes are similar, the positions do not match. This can happen due to various factors, including lack of genomic synteny between *C. lanatus* and *C. amarus* especially with QTL-seq approach which involves alignment with the reference genome of a *C. lanatus* accession. In the current experiment, this can be inferred from the differences in genomic sequence data between the two resistant parents. While PI 482276 had 9.41 GB sequence data, identifying 108157/43322 SNP/InDels, IIHR-82 had only 6.69 GB sequence data with 14750/10957 SNP/InDels. This difference may be attributed to wide variability among these two *C. amarus* accessions in the background of reference *C. lanatus* accession genome used for SNP/InDel analysis.

**Table 3.**
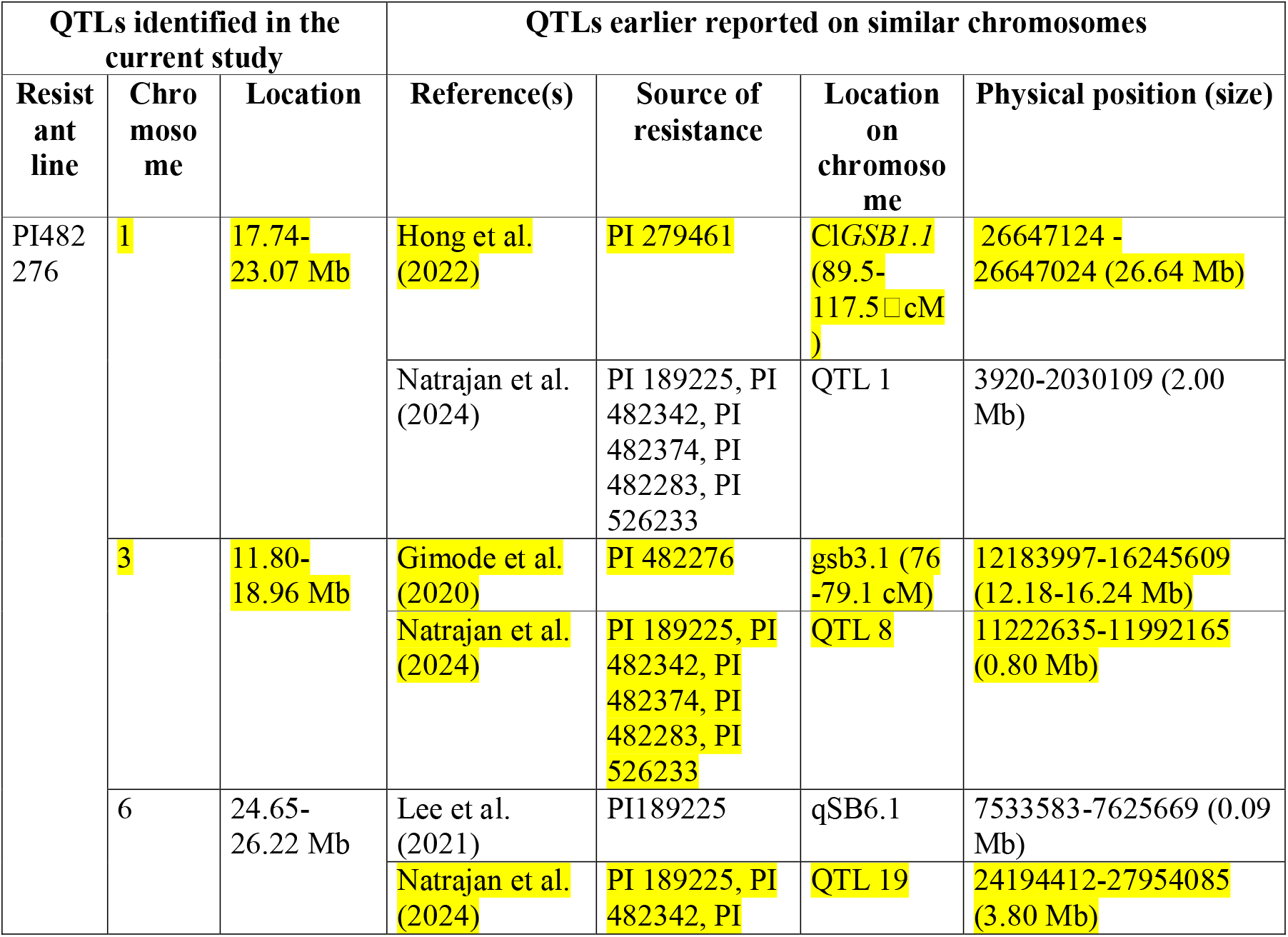
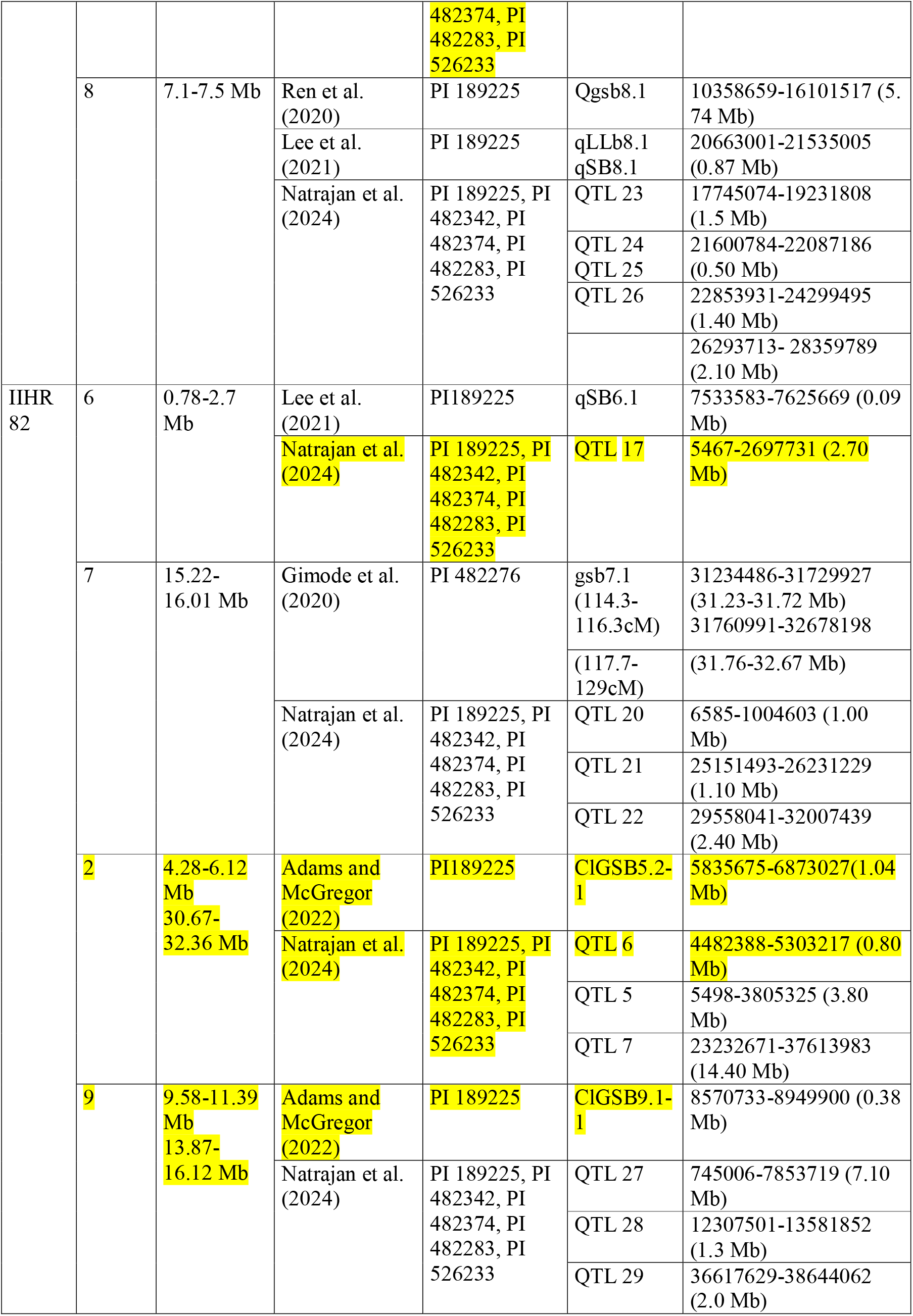
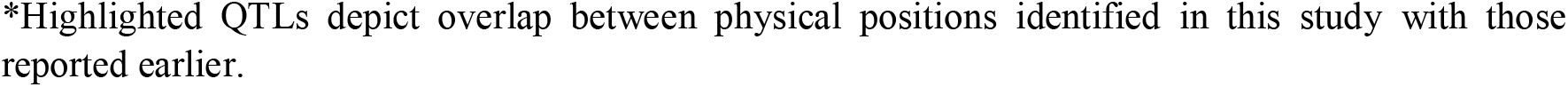
A comparison of the physical position of QTLs for GSB resistance identified in the current study with the previous reports*.

Our results did not match with those of Gimode et al. (2021) except on chromosome 3, although the resistant accession is the same. Similarly, Adams et al. (2022), Lee et al. (2021) and Ren et al. (2020) used the same resistant line (PI 189225) for mapping GSB resistance but obtained a completely different set of QTLs involved in the resistant reaction. Thus, it is interesting to note that no consistency of QTLs was identified across experiments, even with the same resistant parent. Such difference can arise due to the variation in the pathogen strain used in the resistance assessment, suggesting that different loci might control resistance to different species of *Stagonosporopsis* spp. Another possible explanation for this is the use of different rating systems or phenotyping methods. Further, this may also be because of distorted segregation due to wide variation and partial incompatibility observed in the interspecific *C. amarus* × C. *lanatus* crosses (Sandlin et al. 2012, Gimode et al. 2021). This was observed in our experiments as well (Supplementary Table 6).

To conclude, we confirm eight QTLs identified by earlier workers and report two new QTLs using two *Citrullus amarus* accessions. New loci from different sources will allow for pyramiding them into cultivated watermelon for a stronger resistance reaction. However, further efforts have to be made to validate the stability of the QTLs identified in this experiment before deploying them in a resistance breeding program. It is also to be noted that the characterization of the pathogen for probable physiological race differences and the identification of suitable host differentials may provide new insights into the genetics of GSB resistance.

## Compliance with Ethical Standards

### Research involving Human Participants and/or Animals

The authors declare that no human and/or animal participants were involved in this research.

### Informed consent

Not applicable

### Author contribution

Eguru Sreenivasa Rao and DC Lakshmana Reddy conceptualized and supervised the experiments. Sourav Mahapatra and KB Kowsalya performed genetic studies and molecular experiments and prepared the initial draft. Kuldeep Kumar performed the QTL-seq bioinformatics. Saheb Pal and Siddharood Maragal assisted in data cleaning, genomic analysis and contributed to the original draft for corrections. All the authors have read and commented on the manuscript. Sourav Mahapatra, KB Kowsalya and Saheb Pal have contributed equally to the manuscript.

## Acknowledgments

The first author acknowledges ICAR-IARI, New Delhi, India for granting fellowship for PhD programme. The authors thank the Director, ICAR-IIHR for providing the necessary laboratory facilities. The authors are also thankful to SRF’s and YP’s for their constant support throughout different field and laboratory experiments.

## Data Availability

All data generated or analysed during this study are included in this published article and its supplementary information file.

