## Supplementary Tables for "Genetic mapping of *Citrullus amarus* for resistance to gummy stem blight in watermelon"

**Supplementary Table 1: Screening of the parents and the segregating progenies derived from PI 482276 × Arka Manik under artificial inoculation (polyhouse) conditions for gummy stem blight (PDI values as %)**

| **P1** | **P2** | **F_1_** | **F_2-3_** | | **BC1F_1:2_** | **BC2F_1:2_** |
| --- | --- | --- | --- | --- | --- | --- |
| 44.44 | 100.00 | 66.67 | 65.56 | 85.56 | 55.56 | 75.80 |
| 55.56 | 86.67 | 61.90 | 72.23 | 80.74 | 77.51 | 78.33 |
| 77.78 | 87.30 | 70.37 | 68.78 | 68.89 | 74.82 | 71.43 |
| 42.22 | 100.00 | 61.90 | 68.15 | 80.56 | 91.67 | 81.49 |
| 46.67 | 90.48 | 55.56 | 80.42 | 63.10 | 62.96 | 73.15 |
| 44.44 | 85.19 | 64.44 | 66.67 | 91.67 | 71.11 | 44.45 |
| 55.56 | 86.67 | 77.78 | 53.71 | 80.95 | 72.23 | 73.02 |
| 77.78 | 87.30 | 55.56 | 87.94 | 79.63 | 70.37 | 66.67 |
| 42.22 | 100.00 | 55.56 | 72.23 | 65.08 | 55.56 | 83.34 |
| 46.67 | 90.48 | 55.56 | 90.48 | 90.00 | 77.78 | 72.22 |
|  |  | 64.44 | 67.46 | 78.04 | 72.23 | 83.71 |
|  |  | 68.25 | 85.72 | 73.34 | 61.12 | 55.56 |
|  |  | 58.73 | 79.37 | 93.34 | 74.08 | 70.38 |
|  |  | 65.08 | 80.32 | 94.45 | 50.00 | 85.19 |
|  |  | 61.90 | 81.48 | 94.45 | 51.86 | 72.23 |
|  |  | 62.96 | 55.56 | 100.00 | 57.04 | 100.00 |
|  |  | 46.67 | 73.34 | 74.45 | 55.56 | 100.00 |
|  |  | 68.89 | 73.34 | 94.45 | 69.45 | 91.11 |
|  |  | 55.56 | 63.50 | 88.89 | 77.78 | 85.19 |
|  |  | 55.56 | 73.33 | 64.45 | 71.11 | 72.23 |
|  |  | 66.67 | 66.67 | 75.56 | 72.23 | 62.96 |
|  |  | 65.08 | 61.91 | 72.70 | 62.96 | 62.70 |
|  |  | 58.73 | 79.52 | 92.60 | 61.12 | 70.37 |
|  |  | 55.56 | 72.23 | 90.74 | 55.56 |  |
|  |  |  | 88.36 | 91.27 | 62.97 |  |
|  |  |  | 50.00 |  | 55.56 |  |
|  |  |  | 69.45 |  | 66.67 |  |
|  |  |  | 73.34 |  | 70.38 |  |
|  |  |  | 69.26 |  | 55.56 |  |

**Supplementary Table 2:** **Screening of the parents and segregating progenies derived from IIHR 82 × Arka Manik under artificial inoculation (polyhouse) conditions for gummy stem blight (PDI values as %)**

| **P1** | **P2** | **F_1_** | **F_2-3_** | | **BC1F_1:2_** | **BC2F_1:2_** |
| --- | --- | --- | --- | --- | --- | --- |
| 62.96 | 82.22 | 58.73 | 72.22 | 89.42 | 75.56 | 74.08 |
| 67.78 | 83.33 | 55.56 | 48.89 | 91.67 | 79.37 | 77.78 |
| 62.96 | 88.89 | 70.37 | 59.26 | 92.6 | 62.97 | 72.23 |
| 68.25 | 100.00 | 55.56 | 67.59 | 95.93 | 68.89 | 72.23 |
| 65.08 | 92.59 | 55.56 | 74.61 | 88.89 | 79.26 | 83.34 |
| 67.78 | 100.00 | 55.56 | 63.89 | 69.45 | 72.62 | 76.51 |
| 66.67 | 87.30 | 70.37 | 71.75 | 91.11 | 77.78 | 74.08 |
| 55.56 | 100.00 | 70.37 | 61.67 | 90.74 | 65.93 | 80.43 |
| 55.56 | 100.00 | 61.90 | 68.52 | 96.83 | 66.67 | 92.60 |
| 55.56 | 100.00 | 62.96 | 60 | 90.48 | 55.56 | 73.15 |
| 66.67 | 93.65 | 77.78 | 94.45 | 84.45 | 71.30 | 83.34 |
| 68.25 | 100.00 | 82.22 | 91.11 | 94.45 | 75.00 | 92.60 |
| 58.73 | 96.83 | 77.78 | 85.19 | 74.08 | 90.75 | 93.34 |
| 64.44 | 93.65 | 60.00 | 72.7 | 87.84 | 77.78 | 84.45 |
| 63.33 |  | 55.56 | 74.08 | 68.52 | 76.85 | 77.78 |
|  |  | 64.44 | 65 | 64.29 | 70.37 | 91.49 |
|  |  | 77.78 | 77.78 | 75.13 | 55.56 | 82.23 |
|  |  | 72.22 | 57.15 | 77.04 | 85.56 | 79.63 |
|  |  | 55.56 | 81.22 | 93.65 | 90.48 | 80.56 |
|  |  | 74.07 | 74.08 | 83.34 | 61.12 | 80.96 |
|  |  | 77.78 | 80.56 | 70.37 | 55.56 | 77.78 |
|  |  | 60.00 | 77.78 | 80 |  | 79.63 |
|  |  | 66.67 | 74.6 | 79.26 |  | 80.56 |
|  |  | 66.67 | 78.71 | 62.97 |  | 75.19 |
|  |  |  | 82.22 | 72.23 |  | 77.78 |
|  |  |  | 85.45 | 74.08 |  | 80.00 |
|  |  |  | 66.67 | 79.37 |  |  |
|  |  |  | 88.89 | 80.74 |  |  |
|  |  |  | 66.67 | 70.24 |  |  |
|  |  |  | 82.41 | 75.93 |  |  |
|  |  |  | 75.56 | 68.26 |  |  |
|  |  |  | 66.67 | 66.67 |  |  |
|  |  |  | 67.41 | 89.82 |  |  |
|  |  |  | 75 | 77.78 |  |  |
|  |  |  | 82.97 | 82.41 |  |  |
|  |  |  | 86.11 | 80.68 |  |  |
|  |  |  | 86.11 | 88.89 |  |  |
|  |  |  | 79.63 | 88.89 |  |  |
|  |  |  | 89.42 | 100 |  |  |

**Supplementary Table 3. Number of plants/families screened under artificial inoculation (polyhouse) and sick plot (field) conditions for gummy stem blight**

| **Cross** | **Generations** | **Total number of plants/families screened** | |
| --- | --- | --- | --- |
|  |  | Artificial inoculation | Sick plot screening |
| IIHR 82 × Arka Manik | P_1_ (R) | 15 plants | 10 plants |
|  | P_2_ (S) | 14 plants | 10 plants |
|  | F_1_ | 24 plants | 20 plants |
|  | F_2:3_ | 78 families | 78 families |
|  | BC1 F_1:2_ | 21 families | - |
|  | BC2 F_1:2_ | 26 families | - |
| IIHR 617 × Arka Manik | P_1_ (R) | 10 plants | 10 plants |
|  | P_2_ (S) | 10 plants | 10 plants |
|  | F_1_ | 24 plants | 10 plants |
|  | F_2:3_ | 54 families | 54 families |
|  | BC1 F_1:2_ | 29 families | - |
|  | BC2 F_1:2_ | 23 families | - |

14 plants were screened under each family of F_2:3_, BC1F_1-2_ and BC2F_1-2_ under polyhouse and field conditions.

**Supplementary Table 4: Percent Disease Index of the F_2:3_ populations derived from PI 482276 × Arka Manik under sick plot (field) conditions for gummy stem blight**

| **Sl. No.** | **Plant ID** | **PDI (30th DAP)** | **PDI (45 DAP)** | **PDI (60 DAP)** | **PDI (75 DAP)** | **PDI (90 DAP)** |
| --- | --- | --- | --- | --- | --- | --- |
| 1 | 6 | 2.5 (6.46) ± 2.5 | 27.5 (31.59) ± 2.5 | 37.5 (37.66) ± 7.5 | 52.5 (46.42) ± 2.5 | 95 (80.77) ± 5 |
| 2 | 11 | 6.25 (10.35) ± 6.25 | 18.75 (25.34) ± 6.25 | 31.25 (33.87) ± 6.25 | 62.5 (52.48) ± 12.5 | 100 (90) ± 0 |
| 3 | 12 | 7.5 (15.67) ± 2.5 | 32.5 (34.73) ± 2.5 | 37.5 (37.74) ± 2.5 | 60 (50.75) ± 0 | 90 (71.54) ± 0 |
| 4 | 20 | 2.5 (6.46) ± 2.5 | 27.5 (31.59) ± 2.5 | 50 (44.98) ± 5 | 62.5 (52.31) ± 7.5 | 95 (77.0) ± 0 |
| 5 | 26 | 2.5 (6.46) ± 2.5 | 35 (36.05) ± 10 | 50 (44.98) ± 15 | 72.5 (58.37) ± 2.5 | 95 (77.05) ± 0 |
| 6 | 28 | 17.5 (24.21) ± 7.5 | 45 (42.11) ± 0 | 55 (47.87) ± 5 | 60 (50.87) ± 10 | 87.5 (69.36) ± 2.5 |
| 7 | 29 | 2.5 (6.46) ± 2.5 | 30 (33.2) ± 0 | 40 (39.19) ± 5 | 48.5 (44.98) ± 5 | 85 (67.47) ± 5 |
| 8 | 32 | 7.5 (15.67) ± 2.5 | 27.5 (31.59) ± 2.5 | 45 (42.05) ± 10 | 60 (50.87) ± 10 | 95 (80.77) ± 5 |
| 9 | 33 | 15 (21.45) ± 10 | 31.25 (33.87) ± 6.25 | 41.88 (40.3) ± 1.88 | 61.88 (51.92) ± 6.88 | 97.5 (83.52) ± 2.5 |
| 10 | 34 | 12.5 (20.6) ± 2.5 | 30 (33.2) ± 0 | 35 (36.26) ± 0 | 55 (47.87) ± 5 | 85 (67.47) ± 5 |
| 11 | 39 | 10 (18.43) ± 0 | 35 (36.21) ± 5 | 47.5 (43.53) ± 7.5 | 67.5 (55.63) ± 12.5 | 95 (80.77) ± 5 |
| 12 | 40 | 7.5 (15.67) ± 2.5 | 25 (29.99) ± 0 | 37.5 (37.66) ± 7.5 | 67.5 (55.36) ± 7.5 | 90 (71.54) ± 0 |
| 13 | 42 | 2.5 (6.46) ± 2.5 | 27.5 (31.59) ± 2.5 | 32.5 (34.73) ± 2.5 | 55 (47.87) ± 5 | 90 (76.71) ± 10 |
| 14 | 43 | 10 (18.43) ± 0 | 27.5 (31.59) ± 2.5 | 32.5 34.73) ± 2.5 | 50 (44.98) ± 0 | 82.5 (65.3) ± 2.5 |
| 15 | 44 | 5 (9.21) ± 5 | 27.5 (31.59) ± 2.5 | 30 (33.12) ± 5 | 47.5 (43.55) ± 2.5 | 85 (67.47) ± 5 |
| 16 | 48 | 5 (12.92) ± 0 | 35 (36.26) ± 0 | 42.5 (40.67) ± 2.5 | 49.5 (44.87) ± 5 | 82.5 (65.3) ± 2.5 |
| 17 | 50 | 0 (0) ± 0 | 37.5 (37.49) ± 12.5 | 40 (39.09) ± 10 | 60 (50.78) ± 5 | 90 (72.12) ± 5 |
| 18 | 52 | 3.13 (7.24) ± 3.13 | 25 (29.99) ± 0 | 45.63 (42.41) ± 10.63 | 75 (59.98) ± 0 | 100 (90) ± 0 |
| 19 | 56 | 12.5 (19.74) ± 7.5 | 40 (39.22) ± 0 | 40 (39.22) ± 0 | 55 (47.85) ± 0 | 82.5 (65.3) ± 2.5 |
| 20 | 59 | 15.63 (22.23) ± 9.38 | 35 (36.05) ± 10 | 38.13 (38.04) ± 6.88 | 50 (44.98) ± 0 | 96.88 (82.75) ± 3.13 |
| 21 | 60 | 7.5 (11.39) ± 7.5 | 27.5 (31.59) ± 2.5 | 45 (42.05) ± 10 | 65 (53.71) ± 0 | 92.5 (78.59) ± 7.5 |
| 22 | 61 | 2.5 (6.46) ± 2.5 | 27.5 (31.59) ± 2.5 | 30 (33.12) ± 5 | 52.5 (46.42) ± 2.5 | 84.5 (68.36) ± 2.5 |
| 23 | 62 | 5 (9.21) ± 5 | 25 (29.99) ± 0 | 27.5 (31.59) ± 2.5 | 60.5 (51.3) ± 2.5 | 97.5 (83.52) ± 2.5 |
| 24 | 63 | 20 (25.81) ± 10 | 35 (36.05) ± 10 | 40 (39.09) ± 10 | 47.5 (43.5) ± 12.5 | 87.5 (70.23) ± 7.5 |
| 25 | 64 | 0 (0) ± 0 | 30 (33.12) ± 5 | 40 (39.19) ± 5 | 66.5 (57.85) ± 0 | 97.5 (83.52) ± 2.5 |
| 26 | 65 | 5 (12.92) ± 0 | 35 (36.05) ± 10 | 42.5 (40.67) ± 2.5 | 72.5 (58.37) ± 2.5 | 100 (90) ± 0 |
| 27 | 66 | 7.5 (11.39) ± 7.5 | 35 (36.26) ± 0 | 42.5 (40.62) ± 7.5 | 57.5 (49.3) ± 2.5 | 90 (76.71) ± 10 |
| 28 | 67 | 7.5 (15.67) ± 2.5 | 35 (36.21) ± 5 | 45 (42.1) ± 5 | 65 (53.91) ± 10 | 100 (90) ± 0 |
| 29 | 69 | 12.5 (20.6) ± 2.5 | 40 (38.92) ± 15 | 45 (41.97) ± 15 | 56.5 (47.35) ± 7.5 | 85 (67.47) ± 5 |
| 30 | 71 | 5 (9.21) ± 5 | 32.5 (34.6) ± 7.5 | 45 (42.1) ± 5 | 65 (53.76) ± 5 | 100 (90) ± 0 |
| 31 | 72 | 10 (17.85) ± 5 | 40 (39.19) ± 5 | 45 (42.1) ± 5 | 60 (50.78) ± 5 | 90 (76.71) ± 10 |
| 32 | 73 | 18.75 (25.34) ± 6.25 | 46.88 (43.19) ± 3.13 | 46.88 (43.19) ± 3.13 | 55.25 (47.6) ± 6.25 | 78.13 (62.15) ± 3.13 |
| 33 | 74 | 2.5 (6.46) ± 2.5 | 32.5 (34.6) ± 7.5 | 37.5 (37.74) ± 2.5 | 67.5 (55.24) ± 2.5 | 100 (90) ± 0 |
| 34 | 75 | 7.5 (11.39) ± 7.5 | 41.67 (40.12) ± 8.34 | 50.84 (45.47) ± 9.17 | 82.92 (67.17) ± 12.09 | 97.92 (84.09) ± 2.09 |
| 35 | 77 | 6.25 (10.35) ± 6.25 | 25 (29.99) ± 0 | 56.25 (48.6) ± 6.25 | 81.25 (64.62) ± 6.25 | 93.75 (79.63) ± 6.25 |
| 36 | 79 | 14.59 (22.39) ± 2.09 | 25 (29.99) ± 0 | 29.17 (32.62) ± 4.17 | 64.59 (53.67) ± 10.42 | 91.67 (77.94) ± 8.34 |
| 37 | 80 | 7.5 (11.39) ± 7.5 | 27.5 (31.59) ± 2.5 | 40 (39.19) ± 5 | 57.5 (49.35) ± 7.5 | 90 (76.71) ± 10 |
| 38 | 81 | 7.5 (15.67) ± 2.5 | 32.5 (34.73) ± 2.5 | 42.5 (40.52) ± 12.5 | 62.5 (52.48) ± 12.5 | 95 (80.77) ± 5 |
| 39 | 82 | 7.5 (15.67) ± 2.5 | 25 (29.99) ± 0 | 31.25 (33.87) ± 6.25 | 68.75 (59.03) ± 3.75 | 96.5 (84.71) ± 10 |
| 40 | 83 | 4.17 (8.38) ± 4.17 | 30 (33.12) ± 5 | 50.84 (45.47) ± 9.17 | 67.5 (56.08) ± 17.5 | 91.67 (77.94) ± 8.34 |
| 41 | 84 | 12.5 (14.99) ± 12.5 | 50 (44.98) ± 0 | 57.29 (49.17) ± 1.04 | 76.04 (60.93) ± 7.29 | 96.88 (82.75) ± 3.13 |
| 42 | 85 | 10 (13.28) ± 10 | 42.5 (40.62) ± 7.5 | 50 (44.98) ± 5 | 65 (53.76) ± 5 | 100 (90) ± 0 |
| 43 | 86 | 7.5 (15.67) ± 2.5 | 32.5 (34.73) ± 2.5 | 50 (44.98) ± 5 | 57.5 (49.35) ± 7.5 | 85 (67.47) ± 5 |
| 44 | 87 | 2.5 (6.46) ± 2.5 | 27.5 (31.59) ± 2.5 | 42.5 (40.62) ± 7.5 | 66.5 (59.3) ± 2.5 | 97.5 (83.52) ± 2.5 |
| 45 | 88 | 0 (0) ± 0 | 31.25 (33.87) ± 6.25 | 37.5 (37.49) ± 12.5 | 62.5 (52.48) ± 12.5 | 87.5 (74.99) ± 12.5 |
| 46 | 89 | 12.5 (20.6) ± 2.5 | 30 (33.12) ± 5 | 32.5 (34.73) ± 2.5 | 55 (47.87) ± 5 | 80 (63.58) ± 5 |
| 47 | 90 | 2.5 (6.46) ± 2.5 | 24.38 (29.42) ± 5.63 | 27.5 (31.59) ± 2.5 | 50 (44.98) ± 0 | 86.25 (68.23) ± 1.25 |
| 48 | 91 | 15 (22.78) ± 0 | 52.5 (46.42) ± 2.5 | 52.5 (46.42) ± 2.5 | 57.5 (49.3) ± 2.5 | 82.5 (65.3) ± 2.5 |
| 49 | 92 | 6.25 (10.35) ± 6.25 | 29.17 (32.62) ± 4.17 | 41.67 (39.88) ± 16.67 | 62.5 (52.48) ± 12.5 | 90.63 (77.16) ± 9.38 |
| 50 | 93 | 2.5 (6.46) ± 2.5 | 42.5 (40.67) ± 2.5 | 57.5 (49.44) ± 12.5 | 67.5 (55.63) ± 12.5 | 82.5 (65.76) ± 7.5 |
| 51 | 94 | 0 (0) ± 0 | 31.25 (33.87) ± 6.25 | 31.25 (33.87) ± 6.25 | 50 (44.98) ± 0 | 93.75 (79.63) ± 6.25 |
| 52 | 95 | 9.17 (17.6) ± 0.84 | 29.59 (32.94) ± 0.42 | 36.67 (37.23) ± 3.34 | 59.17 (50.26) ± 0.84 | 91.25 (73.16) ± 3.75 |
| 53 | 96 | 7.5 (11.39) ± 7.5 | 35 (36.05) ± 10 | 50 (44.98) ± 20 | 72.5 (58.97) ± 12.5 | 100 (90) ± 0 |
| 54 | 97 | 5 (9.21) ± 5 | 27.5 (31.59) ± 2.5 | 35 (36.26) ± 0 | 52.5 (46.42) ± 2.5 | 92.5 (78.59) ± 7.5 |
| 55 | 99 | 0 ± 0 | 20 (26.38) ± 5 | 25 (29.88) ± 5 | 50 (44.98) ± 0 | 82.5 (65.3) ± 2.5 |
| 56 | IIHR-617 | 3.13 (7.24) ± 3.13 | 7.19 (15.52) ± 0.94 | 17.5 (24.71) ± 1.25 | 27.82 (31.79) ± 2.82 | 63.75 (53.2) ± 11.25 |
| 57 | Arka Manik | 5 (12.92) ± 0 | 41.67 (40.19) ± 0 | 66.67 (54.72) ± 0 | 91.67 (73.2) ± 0 | 100 (90) ± 0 |
| 58 | F_1_ | 0.00 | 8.33 | 27.78 | 50.00 |  |
| 59 | NS 295 | 18.19 (25.23) ± 0.15 | 28.6 (32.32) ± 0.36 | 61.32 (51.52) ± 1.32 | 84.65 (67.15) ± 4.76 | 99.47 (85.95) ± 0.3 |
|  | **F-Calculated** | 1.066 | 2.46 | 1.553 | 1.993 | 1.525 |
|  | **C.D. (at 5%)** | N/A | 8.89 | N/A | 12.20 | N/A |
|  | **C.V.** | 76.08 | 13.00 | 15.32 | 11.86 | 13.22 |
|  | **S.E. (m)±** | 6.79 | 3.13 | 4.31 | 4.30 | 7.13 |

**Supplementary Table. 5. Percent Disease Index of the F_2:3_ population derived from IIHR 82× Arka Manik sick plot (field) conditions for gummy stem blight**

| **Sl. No.** | **Plant ID** | **PDI (30th DAP)** | **PDI (45 DAP)** | **PDI (60 DAP)** | **PDI (75 DAP)** | **PDI (90 DAP)** |
| --- | --- | --- | --- | --- | --- | --- |
| 1 | 1 | 0 (0) ± 0 | 27.5 (31.59) ± 2.5 | 38.75 (38.48) ± 1.25 | 58.75 (50.03) ± 3.75 | 97.5 (83.52) ± 2.5 |
| 2 | 5 | 5 (9.21) ± 5 | 37.5 (37.49) ± 12.5 | 42.5 (40.37) ± 17.5 | 60 (50.75) ± 0 | 85 (73.54) ± 0 |
| 3 | 9 | 0 (0) ± 0 | 32.5 (34.6) ± 7.5 | 47.5 (43.53) ± 7.5 | 72.5 (58.97) ± 12.5 | 100 (90) ± 0 |
| 4 | 10 | 7.5 (11.39) ± 7.5 | 47.5 (43.5) ± 12.5 | 52.5 (46.46) ± 12.5 | 82.5 (65.3) ± 2.5 | 100 (90) ± 0 |
| 5 | 12 | 7.5 (15.67) ± 2.5 | 37.5 (37.49) ± 12.5 | 50 (44.98) ± 0 | 77.5 (61.98) ± 7.5 | 95 (80.77) ± 5 |
| 6 | 13 | 5.63 (13.69) ± 0.63 | 44.38 (41.75) ± 0.63 | 58.13 (49.66) ± 1.88 | 71.88 (57.98) ± 3.13 | 96.875 (82.75) ± 3.13 |
| 7 | 16 | 2.5 (6.46) ± 2.5 | 45 (42.1) ± 5 | 67.5 (55.36) ± 7.5 | 80 (70.37) ± 20 | 97.5 (83.52) ± 2.5 |
| 8 | 18 | 6.67 (14.84) ± 1.67 | 38.34 (38.22) ± 3.34 | 47.5 (43.55) ± 2.5 | 82.5 (65.76) ± 7.5 | 100 (90) ± 0 |
| 9 | 20 | 5 (12.92) ± 0 | 45 (42.05) ± 10 | 77.5 (61.69) ± 2.5 | 92.5 (74.29) ± 2.5 | 100 (90) ± 0 |
| 10 | 24 | 5 (12.92) ± 0 | 55 (47.87) ± 5 | 67.5 (55.36) ± 7.5 | 87.5 (70.23) ± 7.5 | 100 (90) ± 0 |
| 11 | 26 | 10 (13.28) ± 10 | 45 (41.85) ± 20 | 57.5 (49.35) ± 7.5 | 77.5 (63.9) ± 17.5 | 95 (80.77) ± 5 |
| 12 | 27 | 0 (0) ± 0 | 27.5 (31.59) ± 2.5 | 32.5 (34.6) ± 7.5 | 47.5 (43.55) ± 2.5 | 82.5 (65.76) ± 7.5 |
| 13 | 28 | 2.5 (6.46) ± 2.5 | 38.75 (38.48) ± 1.25 | 50 (44.98) ± 0 | 73.75 (60.01) ± 13.75 | 100 (90) ± 0 |
| 14 | 31 | 0 (0) ± 0 | 31.25 (33.87) ± 6.25 | 35.42 (36.5) ± 2.09 | 58.34 (49.85) ± 8.34 | 100 (90) ± 0 |
| 15 | 32 | 3.13 (7.24) ± 3.13 | 29.17 (32.62) ± 4.17 | 44.79 (41.88) ± 13.54 | 65.63 (54.65) ± 15.63 | 91.665 (77.94) ± 8.34 |
| 16 | 35 | 27.5 (31) ± 12.5 | 62.5 (52.48) ± 12.5 | 72.5 (58.56) ± 7.5 | 80 (63.58) ± 5 | 97.5 (83.52) ± 2.5 |
| 17 | 36 | 7.5 (15.67) ± 2.5 | 27.5 (31.59) ± 2.5 | 45 (42.1) ± 5 | 57.5 (49.35) ± 7.5 | 92.5 (78.59) ± 7.5 |
| 18 | 37 | 12.5 (19.74) ± 7.5 | 40 (39.09) ± 10 | 60 (50.75) ± 0 | 75 (59.98) ± 0 | 95 (80.77) ± 5 |
| 19 | 38 | 0 (0) ± 0 | 25 (29.99) ± 0 | 50 (44.98) ± 0 | 50 (44.98) ± 0 | 75 (59.98) ± 0 |
| 20 | 39 | 5 (12.92) ± 0 | 45 (42.11) ± 0 | 50 (44.98) ± 0 | 62.5 (52.23) ± 2.5 | 90 (72.12) ± 5 |
| 21 | 40 | 7.5 (15.67) ± 2.5 | 27.5 (31.59) ± 2.5 | 45 (42.05) ± 10 | 55 (47.87) ± 5 | 95 (80.77) ± 5 |
| 22 | 41 | 5 (9.21) ± 5 | 46.25 (42.71) ± 16.25 | 54.38 (47.6) ± 14.38 | 73.13 (59.01) ± 8.13 | 97.5 (83.52) ± 2.5 |
| 23 | 42 | 7.5 (11.39) ± 7.5 | 52.5 (46.43) ± 7.5 | 57.5 (49.35) ± 7.5 | 67.5 (55.63) ± 12.5 | 97.5 (83.52) ± 2.5 |
| 24 | 43 | 3.13 (7.24) ± 3.13 | 27.5 (31.59) ± 2.5 | 33.13 (35.12) ± 1.88 | 52.5 (46.42) ± 2.5 | 85 (73.77) ± 5 |
| 25 | 45 | 15.42 (22.79) ± 5.42 | 42.5 (40.62) ± 7.5 | 53.75 (47.17) ± 8.75 | 81.25 (64.62) ± 6.25 | 95 (80.77) ± 5 |
| 26 | 46 | 0 (0) ± 0 | 45.84 (42.59) ± 4.17 | 50 (44.98) ± 0 | 76.67 (61.32) ± 6.67 | 95 (80.77) ± 5 |
| 27 | 47 | 12.5 (20.6) ± 2.5 | 42.5 (40.52) ± 12.5 | 45 (42.05) ± 10 | 57.5 (49.35) ± 7.5 | 95 (80.77) ± 5 |
| 28 | 48 | 0 (0) ± 0 | 57.5 (49.6) ± 17.5 | 65 (54.2) ± 15 | 72.5 (59.69) ± 17.5 | 95 (80.77) ± 5 |
| 29 | 49 | 0 (0) ± 0 | 50 (44.98) ± 0 | 50 (44.98) ± 0 | 75 (59.98) ± 0 | 100 (90) ± 0 |
| 30 | 51 | 0 (0) ± 0 | 42.5 (40.52) ± 12.5 | 50 (44.98) ± 5 | 65 (53.76) ± 5 | 97.5 (83.52) ± 2.5 |
| 31 | 52 | 10 (13.28) ± 10 | 40 (38.92) ± 15 | 50 (44.98) ± 15 | 59.13 (50.87) ± 10 | 85 (67.19) ± 0 |
| 32 | 53 | 12.5 (14.99) ± 12.5 | 40 (39.09) ± 10 | 46.25 (42.71) ± 16.25 | 56.25 (48.6) ± 6.25 | 97.5 (83.52) ± 2.5 |
| 33 | 55 | 3.13 (7.24) ± 3.13 | 40.63 (39.48) ± 9.38 | 46.88 (43.19) ± 3.13 | 84.38 (67.73) ± 9.38 | 100 (90) ± 0 |
| 34 | 57 | 10 (17.85) ± 5 | 35 (36.05) ± 10 | 42.5 (40.67) ± 2.5 | 52.5 (46.42) ± 2.5 | 92.5 (74.29) ± 2.5 |
| 35 | 58 | 18.75 (18.87) ± 18.75 | 53.75 (47.17) ± 8.75 | 60 (51.05) ± 15 | 85.63 (67.93) ± 4.38 | 100 (90) ± 0 |
| 36 | 60 | 10 (17.85) ± 5 | 37.5 (37.74) ± 2.5 | 47.5 (43.53) ± 7.5 | 60 (50.75) ± 0 | 90 (71.54) ± 0 |
| 37 | 62 | 8.75 (16.81) ± 3.75 | 48.75 (44.25) ± 11.25 | 59.59 (50.54) ± 5.42 | 73.34 (59.06) ± 6.67 | 95.835 (81.6) ± 4.17 |
| 38 | 64 | 14.38 (22.04) ± 4.38 | 28.13 (31.98) ± 3.13 | 51.88 (48.3) ± 1.88 | 73.63 (59.53) ± 10.63 | 97.5 (83.52) ± 2.5 |
| 39 | 65 | 4.17 (8.38) ± 4.17 | 29.17 (32.62) ± 4.17 | 45 (42.1) ± 5 | 54.17 (47.38) ± 4.17 | 86.25 (68.23) ± 1.25 |
| 40 | 70 | 5 (9.21) ± 5 | 47.5 (43.45) ± 17.5 | 60 (50.87) ± 10 | 75 (60.09) ± 5 | 95 (80.77) ± 5 |
| 41 | 72 | 2.5 (6.46) ± 2.5 | 42.5 (40.62) ± 7.5 | 57.5 (49.35) ± 7.5 | 67.5 (55.36) ± 7.5 | 95 (80.77) ± 5 |
| 42 | 76 | 2.5 (6.46) ± 2.5 | 42.5 (40.67) ± 2.5 | 59.5 (51.3) ± 2.5 | 74.39 (59.08) ± 10 | 90 (72.12) ± 5 |
| 43 | 77 | 2.5 (6.46) ± 2.5 | 32.5 (34.6) ± 7.5 | 32.5 (34.6) ± 7.5 | 50 (44.98) ± 0 | 84.165 (66.53) ± 0.84 |
| 44 | 78 | 0 (0) ± 0 | 31.67 (34.22) ± 1.67 | 40.84 (39.7) ± 0.84 | 56.67 (48.81) ± 1.67 | 75 (59.98) ± 0 |
| 45 | 79 | 23.13 (28.38) ± 8.13 | 50.63 (45.34) ± 5.63 | 53.75 (47.17) ± 8.75 | 75 (59.98) ± 0 | 100 (90) ± 0 |
| 46 | 80 | 0 (0) ± 0 | 65.63 (54.27) ± 9.38 | 70.63 (57.88) ± 14.38 | 79.75 (64.63) ± 16.25 | 93.75 (79.63) ± 6.25 |
| 47 | 82 | 0 (0) ± 0 | 30 (33.12) ± 5 | 52.5 (46.42) ± 2.5 | 65 (53.76) ± 5 | 88.75 (70.4) ± 1.25 |
| 48 | 83 | 23.34 (26.55) ± 18.34 | 60.84 (51.28) ± 5.84 | 60.84 (51.28) ± 5.84 | 80 (63.58) ± 5 | 100 (90) ± 0 |
| 49 | 84 | 10 (17.85) ± 5 | 35 (36.05) ± 10 | 40 (39.09) ± 10 | 57.5 (49.3) ± 2.5 | 85 (67.19) ± 0 |
| 50 | 85 | 7.5 (15.67) ± 2.5 | 37.5 (37.74) ± 2.5 | 50 (44.98) ± 10 | 64.5 (51.63) ± 12.5 | 82.5 (70.29) ± 2.5 |
| 51 | 90 | 12.5 (14.99) ± 12.5 | 50 (44.98) ± 5 | 57.5 (49.35) ± 7.5 | 57.5 (49.44) ± 12.5 | 82.5 (65.3) ± 2.5 |
| 52 | 92 | 2.5 (6.46) ± 2.5 | 45 (42.1) ± 5 | 72.5 (58.37) ± 2.5 | 91.25 (73.16) ± 3.75 | 100 (90) ± 0 |
| 53 | 93 | 0 (0) ± 0 | 45.84 (42.35) ± 20.84 | 67.09 (55.34) ± 12.09 | 87.92 (70.8) ± 7.92 | 100 (90) ± 0 |
| 54 | 94 | 3.13 (7.24) ± 3.13 | 36.25 (37) ± 1.25 | 55 (47.87) ± 5 | 65 (54.2) ± 15 | 96.875 (82.75) ± 3.13 |
| 55 | 95 | 0 (0) ± 0 | 32.5 (34.73) ± 2.5 | 55 (47.85) ± 0 | 60 (50.75) ± 0 | 90 (72.12) ± 5 |
| 56 | 96 | 20 (25.81) ± 10 | 37.5 (37.49) ± 12.5 | 55 (47.87) ± 5 | 72.5 (58.37) ± 2.5 | 97.5 (83.52) ± 2.5 |
| 57 | 97 | 0 (0) ± 0 | 45 (42.1) ± 5 | 65 (53.71) ± 0 | 70 (56.84) ± 5 | 87.5 (69.36) ± 2.5 |
| 58 | 98 | 5.63 (13.69) ± 0.63 | 40 (39.09) ± 10 | 48.13 (43.89) ± 8.13 | 71.25 (58.56) ± 16.25 | 86.87 (69.45) ± 6.88 |
| 59 | 100 | 3.13 (7.24) ± 3.13 | 35.63 (36.6) ± 4.38 | 52.5 (46.42) ± 2.5 | 86.88 (69.45) ± 6.88 | 96.87 (82.75) ± 3.13 |
| 60 | 101 | 7.5 (15.67) ± 2.5 | 52.5 (46.43) ± 7.5 | 62.5 (52.31) ± 7.5 | 82.5 (66.91) ± 12.5 | 100 (90) ± 0 |
| 61 | 102 | 0 (0) ± 0 | 47.5 (43.5) ± 12.5 | 70 (56.77) ± 0 | 90 (72.12) ± 5 | 100 (90) ± 0 |
| 62 | 105 | 0 (0) ± 0 | 50 (44.98) ± 0 | 68.75 (57.13) ± 18.75 | 81.25 (64.62) ± 6.25 | 85.41 (67.57) ± 2.09 |
| 63 | 106 | 2.5 (6.46) ± 2.5 | 48.75 (44.24) ± 13.75 | 56.25 (48.6) ± 6.25 | 77.5 (61.69) ± 2.5 | 100 (90) ± 0 |
| 64 | 108 | 0 (0) ± 0 | 45 (42.05) ± 10 | 50 (44.98) ± 5 | 57.5 (49.3) ± 2.5 | 92.5 (74.29) ± 2.5 |
| 65 | 112 | 6.25 (10.35) ± 6.25 | 60.42 (51) ± 2.09 | 60.42 (51) ± 2.09 | 67.71 (55.35) ± 1.04 | 100 (90) ± 0 |
| 66 | 113 | 0 (0) ± 0 | 55 (47.87) ± 5 | 60 (50.87) ± 10 | 62.5 (52.48) ± 12.5 | 90 (76.7) ± 10 |
| 67 | 115 | 2.09 (5.89) ± 2.09 | 30 (33.12) ± 5 | 47.92 (43.79) ± 2.09 | 77.09 (61.4) ± 2.09 | 100 (90) ± 0 |
| 68 | 116 | 12.5 (20.6) ± 2.5 | 37.5 (37.74) ± 2.5 | 45 (42.1) ± 5 | 57.5 (49.3) ± 2.5 | 97.5 (83.52) ± 2.5 |
| 69 | 117 | 0 (0) ± 0 | 25 (29.99) ± 0 | 37.5 (37.49) ± 12.5 | 50 (44.98) ± 0 | 88.42 (76.36) ± 5 |
| 70 | 118 | 6.25 (10.35) ± 6.25 | 36.88 (37.3) ± 6.88 | 51.25 (45.72) ± 11.25 | 81.88 (66.13) ± 11.88 | 100 (90) ± 0 |
| 71 | 119 | 0 (0) ± 0 | 69.17 (56.86) ± 14.17 | 69.17 (56.86) ± 14.17 | 85.84 (68.3) ± 5.84 | 100 (90) ± 0 |
| 72 | 121 | 0 (0) ± 0 | 42.5 (40.62) ± 7.5 | 65 (53.76) ± 5 | 92.5 (74.29) ± 2.5 | 100 (90) ± 0 |
| 73 | 123 | 10 (17.85) ± 5 | 40 (39.22) ± 0 | 67.5 (59.35) ± 7.5 | 72.5 (61.23) ± 2.5 | 100 (90) ± 0 |
| 74 | 125 | 0 (0) ± 0 | 62.5 (52.22) ± 0 | 75 (59.98) ± 0 | 81.25 (64.32) ± 0 | 100 (90) ± 0 |
| 75 | 127 | 0 (0) ± 0 | 25 (29.99) ± 0 | 37.5 (37.49) ± 12.5 | 75 (67.49) ± 25 | 100 (90) ± 0 |
| 76 | 128 | 3.13 (7.24) ± 3.13 | 34.38 (35.69) ± 9.38 | 56.25 (48.86) ± 18.75 | 68.75 (57.13) ± 18.75 | 100 (90) ± 0 |
| 77 | IIHR 82 | 2.29 (6.18) ± 2.29 | 6.46 (14.72) ± 0.21 | 23.75 (29.15) ± 1.25 | 44.59 (41.86) ± 5.42 | 68.54 (55.97) ± 6.46 |
| 78 | Arka Manik | 5 (12.92) ± 0 | 41.67 (40.19) ± 0 | 66.67 (54.72) ± 0 | 91.67 (73.2) ± 0 | 100 (90) ± 0 |
| 79 | F_1_ | 0.00 | 26.81 | 50.69 | 57.08 | 66.75 |
| 80 | NS 295 | 14.94 (22.73) ± 0.56 | 28 (31.93) ± 0.5 | 61.65 (51.72) ± 0.98 | 85.23 (67.96) ± 6.9 | 99.555 (86.15) ± 0.06 |
|  | **F-Calculated** | 1.656 | 1.941 | 1.717 | 1.707 | 2.77 |
|  | **C.D. (at 5%)** | 18.239 | 13.691 | 14.04 | 17.851 | 15.186 |
|  | **C.V.** | 95.924 | 17.288 | 15.002 | 15.46 | 9.397 |
|  | **S.E. (m)±** | 6.464 | 4.852 | 4.976 | 6.327 | 5.382 |

**Supplementary Table 6: Details of sequencing data generated in the current experiment.**

| **Sample** | **IIHR 82** | **IIHR 82 Resistant Bulk** | **IIHR 82 Susceptible Bulk** | **PI 482276** | **PI 482276 Resistant Bulk** | **PI 482276 Susceptible Bulk** |
| --- | --- | --- | --- | --- | --- | --- |
| **Total number of bases** | 3925380450 | 3346191300 | 3528868200 | 3374754000 | 4702525500 | 3930712500 |
| **Total number of reads** | 26169203 | 22307942 | 23525788 | 22498360 | 31350170 | 26204750 |
| **% bases >=Q20** | 97.43 | 97.57 | 97.54 | 97.47 | 97.57 | 97.43 |
| **% bases >=Q30** | 92.93 | 93.17 | 93.14 | 93.04 | 93.25 | 92.92 |
| **Number of base A** | 1276628262 | 1081986587 | 1136584602 | 1093500908 | 1533424331 | 1285259784 |
| **Number of base T** | 1261815892 | 1069537204 | 1121152682 | 1082496062 | 1518585997 | 1270766381 |
| **Number of base G** | 694321953 | 598278692 | 636969628 | 599981008 | 827341872 | 687795879 |
| **Number of base C** | 692582375 | 596361554 | 634132570 | 598748360 | 823130249 | 686858449 |
| **GC content %** | 35.33 | 35.70 | 36.02 | 35.52 | 35.10 | 34.97 |

**Supplementary Table 7A: Crossability in the *Citrullus amarus* × *Citrullus lanatus* crosses.**

| **Sl. No.** | **Hybrids** | **No. of bold seeds** | **No. of chaffy seeds** | **Germination %** |
| --- | --- | --- | --- | --- |
| 1 | IIHR-82 × IIHR-30 | 458 | 9 | 43.75 |
| 2 | IIHR-82 × IIHR-260 | 438 | 1 | 15.71 |
| 3 | IIHR-82 × IIHR-392 | 324 | 1 | 10.7 |
| 4 | IIHR-82 × IIHR-368 | 280 | 8 | 18.3 |
| 5 | IIHR-82 × IIHR-177 | 604 | 1 | 41.66 |
| 6 | IIHR-82 × IIHR-164 | 309 | 3 | 29.78 |

| **Sl. No.** | **Hybrids** | **No. of bold seeds** | **No. of chaffy seeds** | **Germination %** |
| --- | --- | --- | --- | --- |
| 1 | IIHR-617 × IIHR-271 | 302 | 1 | 15 |
| 2 | IIHR-617 × IIHR-351 | 220 | 1 | 75 |
| 3 | IIHR-617 × IIHR-517 | 155 | 4 | 61.42 |
| 4 | IIHR-617 × IIHR-177 | 215 | 1 | 13.51 |

**Supplementary Table 7B: Crossability in the reciprocal crosses i.e., *Citrullus lanatus* × *Citrullus amarus***

| **Sl. No.** | **Hybrids** | **No. of bold seeds** | **No. of chaffy seeds** | **Germination %** |
| --- | --- | --- | --- | --- |
| 1 | IIHR-30 × IIHR-82 | 254 | 230 | 8 |
| 2 | IIHR-260 × IIHR-82 | 70 | 78 | 12 |
| 3 | IIHR-392 × IIHR-82 | 294 | 12 | 28 |
| 4 | IIHR-368 × IIHR-82 | 380 | 12 | 78 |
| 5 | IIHR-177 × IIHR-82 | 144 | 147 | 32 |
| 6 | IIHR-164 × IIHR-82 | 100 | 205 | 3 |

| **Sl. No.** | **Hybrids** | **No. of bold seeds** | **No. of chaffy seeds** | **Germination %** |
| --- | --- | --- | --- | --- |
| 1 | IIHR-271× IIHR-617 | 327 | 150 | 20 |
| 2 | IIHR-351× IIHR-617 | 131 | 57 | 2 |
| 3 | IIHR-517× IIHR-617 | 40 | 190 | 1 |
| 4 | IIHR-177× IIHR-617 | 170 | 10 | 80 |

**Supplementary Table 8: A comparison of the physical position of QTLs for GSB resistance identified in the current study with the previous reports***

| **QTLs identified in the current study** | | | **QTLs reported earlier on similar chromosomes** | | | |
| --- | --- | --- | --- | --- | --- | --- |
| **Resistant line** | **Chromosome** | **Location** | **Reference(s)** | **Source of resistance** | **QTL name and position** | **Physical position (Size)** |
| PI482276 | 1 | 17.74-23.07 Mb | Hong et al. (2022) | PI 279461 | Cl*GSB1.1* (89.5-117.5 cM) | 26647124 -26647024 (100 bb) |
|  |  |  | Natrajan et al. (2024) | PI 189225, PI 482342, PI 482374, PI 482283, PI 526233 | QTL 1 | 3920-2030109 (2.00 Mb) |
|  | 3 | 11.80-18.96 Mb | Gimode et al. (2020) | PI 482276 | *gsb3.1* (76 -79.1 cM) | 12183997-16245609 (12.18-16.24 Mb) |
|  |  |  | Natrajan et al. (2024) | PI 189225, PI 482342, PI 482374, PI 482283, PI 526233 | QTL 8 | 11222635-11992165 (0.80 Mb) |
|  | 6 | 24.65-26.22 Mb | Lee et al. (2021) | PI189225 | *qSB6.1* | 7533583-7625669 (0.09 Mb) |
|  |  |  | Natrajan et al. (2024) | PI 189225, PI 482342, PI 482374, PI 482283, PI 526233 | QTL 19 | 24194412-27954085 (3.80 Mb) |
|  | 8 | 7.1-7.5 Mb | Ren et al. (2020) | PI 189225 | *Qgsb8.1* | 10358659-16101517 (5. 74 Mb) |
|  |  |  | Lee et al. (2021) | PI 189225 | *qLLb8.1*  *qSB8.1* | 20663001-21535005 (0.87 Mb) |
|  |  |  | Natrajan et al. (2024) | PI 189225, PI 482342, PI 482374, PI 482283, PI 526233 | QTL 23 | 17745074-19231808 (1.5 Mb)  21600784-22087186 (0.50 Mb) |
|  |  |  |  |  | QTL 24 | 21600784-22087186 (0.50 Mb) |
|  |  |  |  |  | QTL 25 | 22853931-24299495 (1.40 Mb) |
|  |  |  |  |  | QTL 26 | 26293713- 28359789 (2.10 Mb) |
| IIHR 82 | 6 | 0.78-2.7 Mb | Lee et al. (2021) | PI189225 | *qSB6.1* | 7533583-7625669 (0.09 Mb) |
|  |  |  | Natrajan et al. (2024) | PI 189225, PI 482342, PI 482374, PI 482283, PI 526233 | QTL 17 | 5467-2697731 (2.70 Mb) |
|  | 7 | 15.22-16.01 Mb | Gimode et al. (2020) | PI 482276 | *gsb7.1* (114.3-116.3 cM) | 31234486-31729927 (31.23-31.72 Mb) |
|  |  |  |  |  | (117.7-129 cM) | 31760991-32678198 (31.76-32.67 Mb) |
|  |  |  | Natrajan et al. (2024) | PI 189225, PI 482342, PI 482374, PI 482283, PI 526233 | QTL 20 | 6585-1004603 (1.00 Mb) |
|  |  |  |  |  | QTL 21 | 25151493-26231229 (1.10 Mb) |
|  |  |  |  |  | QTL 22 | 29558041-32007439 (2.40 Mb) |
|  | 2 | 4.28-6.12 Mb  30.67-32.36 Mb | Adams and McGregor (2022) | PI189225 | *ClGSB5.2-1* | 5835675-6873027(1.04 Mb) |
|  |  |  | Natrajan et al. (2024) | PI 189225, PI 482342, PI 482374, PI 482283, PI 526233 | QTL 6 | 4482388-5303217 (0.80 Mb) |
|  |  |  |  |  | QTL 5 | 5498-3805325 (3.80 Mb) |
|  |  |  |  |  | QTL 7 | 23232671-37613983 (14.40 Mb) |
|  | 9 | 9.58-11.39 Mb  13.87-16.12 Mb | Adams and McGregor (2022) | PI 189225 | ClGSB9.1-1 | 8570733-8949900 (0.38 Mb) |
|  |  |  | Natrajan et al. (2024) | PI 189225, PI 482342, PI 482374, PI 482283, PI 526233 | QTL 27 | 745006-7853719 (7.10 Mb) |
|  |  |  |  |  | QTL 28 | 12307501-13581852 (1.3 Mb) |
|  |  |  |  |  | QTL 29 | 36617629-38644062 (2.0 Mb) |

*Highlighted QTLs depict overlap between physical positions identified in this study with those reported earlier.
